# Lipid nanoemulsion incorporating DOTAP reverse micelles as clinically translatable carriers of ALDH inhibitors for lung cancer therapy

**DOI:** 10.1101/2025.06.16.659868

**Authors:** Maria Irujo, Laura Poussereau, Maïssam Ezziani, Clotilde Joubert, Chloé Prunier, Julien Vollaire, Véronique Josserand, Cédric Sarazin, Axel Kattar, Daphna Fenel, Eleftherios Zarkadas, Alice Gaudin, Mileidys Perez, Isabelle Texier

**Author notes:** **Corresponding authors** Isabelle Texier,; Alice Gaudin.

## Abstract

Lung cancer remains the most prevalent malignancy worldwide and the leading cause of cancer-related deaths. In this study, we designed and optimized a lipid nanosystem incorporating the cationic lipid DOTAP by utilizing reverse micelle structures to enhance pulmonary tropism. Fluorescence quenching assays confirmed the incorporation of reverse micelles into the oily core, thus validating the structural integrity of the system. This nanosystem was tailored for the encapsulation of ABD0171, a potent inhibitor of ALDH1A3, an enzyme strongly associated with chemoresistance in lung cancer. Physicochemical characterization revealed robust colloidal properties with a particle diameter of 60 nm, a surface charge of +45 mV, and a drug encapsulation yield of 99%. *In vitro*, formulations with/without DOTAP demonstrated high efficacy against epithelial-like H358 cells derived from human bronchioalveolar carcinoma, with IC50 values < 5 µM. Chicken ChorioAllantoic Membrane (CAM) evaluations demonstrated good tolerability of the DOTAP-containing formulation and significant H358 tumor reduction by 28% compared with the untreated control, while maintaining a favorable safety profile. By combining industrially feasible methods with FDA-approved components, this study demonstrated the potential of DOTAP-containing lipid nanosystems as scalable platforms for the delivery of ALDH inhibitors for lung cancer treatment. These findings provide a pathway for future applications in cancer therapy, bridging the gap between nanosystem innovation and clinical translation.

Graphical abstract

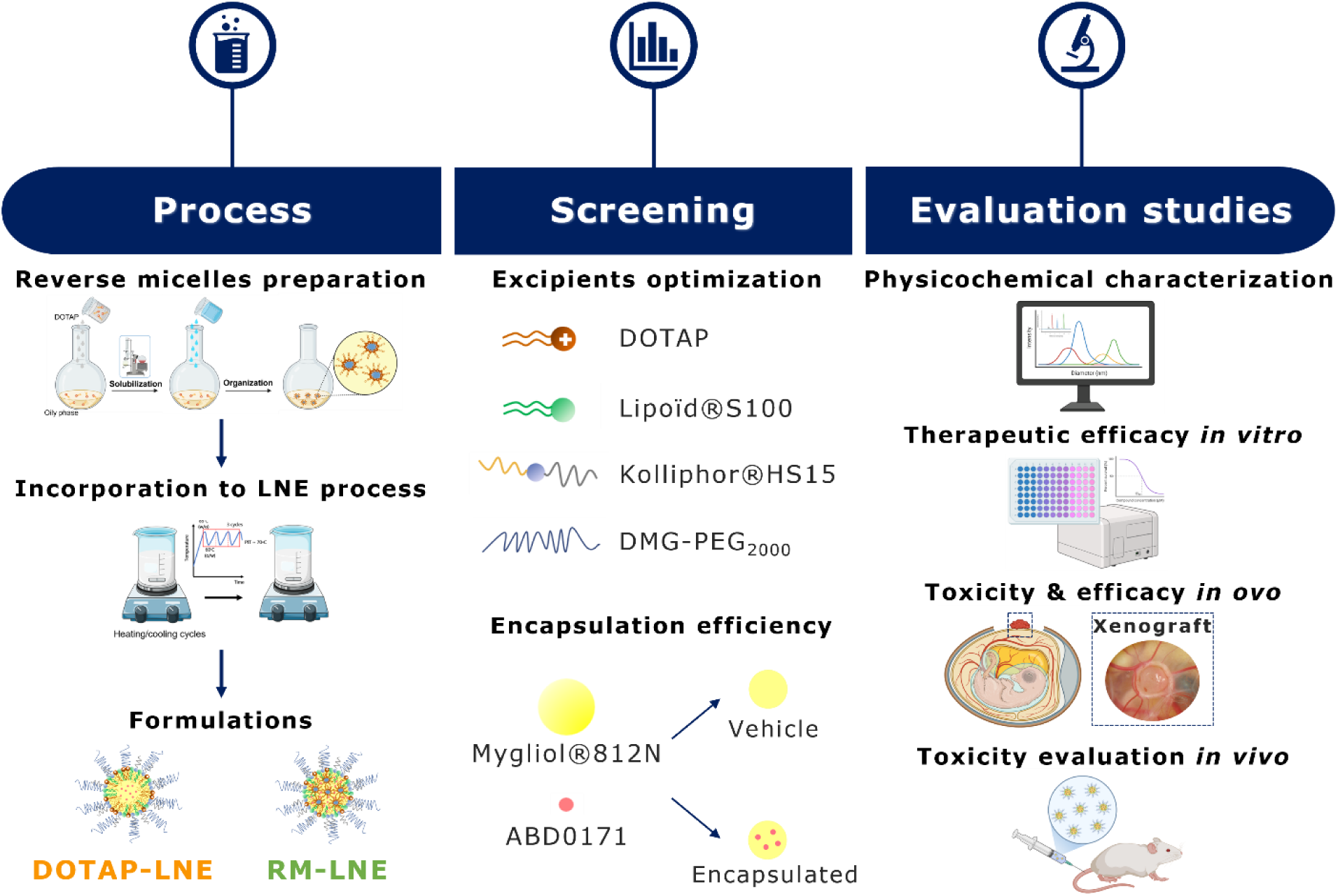

## INTRODUCTION

Lung cancer remains the most prevalent malignancy worldwide and is the leading cause of cancer-related deaths with 2.2 million new cases diagnosed annually (1,2). Non-small cell lung cancer (NSCLC) accounts for approximately 80–85% of all lung cancer cases and includes adenocarcinoma, squamous cell carcinoma, and large cell carcinoma as the main histological subtypes (3). Advances in screening techniques, such as low-dose computed tomography (LDCT), have improved early detection of the disease; however, most patients are still diagnosed at advanced stages, leading to poor prognosis (3,4). As a result, the five-year survival rate for all combined stages remains as low as 28%, emphasizing the critical need for innovative therapeutic strategies (1,3). Current therapeutic options for NSCLC include surgery, chemotherapy, radiotherapy, and targeted therapy. While targeted therapies and immunotherapies have demonstrated promising results, resistance to these treatments, as well as side effects, frequently emerge, driven by tumor heterogeneity and adaptive mechanisms, which significantly hamper their long-term effectiveness (1,2,5). Consequently, there is an urgent need to develop novel therapeutic approaches to address these challenges and to improve patient outcomes.

One promising target in NSCLC therapy is the class 1 aldehyde dehydrogenase (ALDH) subfamily of enzymes, particularly the ALDH1A1 and ALDH1A3 isoforms, which have been shown to be implicated in tumorigenicity, chemoresistance, and disease progression (6–8). ALDH1A3 has been identified as a biomarker of cancer stem-like cells and is associated with an enhanced tumor-initiating potential. Targeting ALDH1A3 sensitizes NSCLC cells to existing treatments such as chemotherapy and radiotherapy, potentially by amplifying reactive oxygen species (ROS) production (6). However, the clinical translation of ALDH inhibitors (ALDHin) remains limited by their poor solubility, short plasma half-life, and rapid clearance *in vivo* (7). This calls for the development of advanced nanocarrier systems to enhance ALDHin delivery and therapeutic use.

Lipid nanocarriers, including lipid nanoparticles (LNPs), lipid nanocapsules (LNCs), and lipid nanoemulsions (LNEs), have emerged as promising platforms for drug delivery. Their ability to encapsulate hydrophobic molecules, combined with their biocompatibility and potential for targeted delivery, make them ideal candidates for cancer therapy (9–11). Despite the major advances in the development of such nanosystems over the last decades, ensuring efficient delivery to the desired therapeutic site remains a critical challenge. Passive targeting, which relies on the enhanced permeability and retention (EPR) effect, often fails to bypass rapid clearance by the reticuloendothelial system (RES), significantly reducing circulation time (12). Active targeting, on the other hand, involves functionalizing nanoparticle surfaces with specific ligands, and has shown promise in preclinical studies (11). However, its large-scale translation is hindered by high production costs and limited clinical benefits (13). An innovative approach to nanoparticle engineering is endogenous targeting, which exploits the interactions between the nanoparticle surface and plasma proteins for direct organ-specific delivery (14). In particular, the incorporation of cationic lipids, such as DOTAP (1,2-dioleoyl-3-trimethylammonium-propane) into nanoparticle formulations has been shown to enhance lung tropism through a three-step mechanism: (a) desorption of polyethylene glycol (PEG) chains from the nanoparticle surface, (b) adsorption of plasma proteins onto the DOTAP-modified surface, and (c) subsequent recognition of these protein-nanoparticle complexes by adhesion molecule receptors specifically expressed in lung tissue (15). Building upon this mechanism, and with a focus on clinical translation, our study explores the incorporation of DOTAP into lipid nanoemulsions as a strategy to facilitate pulmonary delivery of ABD0171, a potent and selective ALDH1A3 inhibitor (16).

DOTAP is commonly used in lipid nanoparticles (LNPs) for mRNA delivery, where it interacts electrostatically with the negatively charged mRNA. This process is typically achieved through microfluidic mixing, which enables spontaneous self-assembly of the nanoparticles (15). In that context, it has been suggested that the cationic lipid DOTAP must be located within the core of the nanoparticle to enable the three-step mechanism required for efficient delivery and selective organ targeting (15). However, in our case, the therapeutic compound is a lipophilic ALDH inhibitor with no permanent charge, and thus cannot be incorporated via electrostatic complexation as in mRNA–LNP systems. To address this, we adapted a strategy previously described in LNCs, where surfactants such as AOT have been used to form reverse micelles and enable encapsulation of hydrophilic drugs within the oily core (17). Building on this concept, we designed a nanoemulsion incorporating DOTAP reverse micelles, not to deliver active compound, but to integrate the targeting lipid (DOTAP) into the core structure of the nanocarrier. This strategy enables controlled incorporation of DOTAP within the oil core, potentially favoring targeting mechanisms similar to those observed in DOTAP-mRNA–LNPs. However, from a clinical translation perspective, the addition of new components such as DOTAP, and the design of more complex internal architectures like reverse micelles, may raise formulation and regulatory challenges. Their use must therefore be clearly justified by a demonstrated therapeutic benefit and manufacturing feasibility.

In this study, we particularly developed, optimized, and characterized a novel lipid nanoemulsion formulation incorporating DOTAP through reverse micelle structures (RM-LNE), constituting a stable and efficient colloidal drug delivery system capable of delivering ABD0171 to pulmonary cells, to unlock its therapeutic potential. We assessed and compared the physicochemical properties and therapeutic efficacy of three ABD0171-loaded formulations **(Figure 1)**: 1) LNE incorporating DMPE-PEG_2000_ is a PEGylated version of the nanoemulsion developed by Heurtault et al. (18), 2) DOTAP-LNE is the same formulation incorporating DOTAP by directly mixing it with the other LNE components, and 3) RM-LNE incorporates DOTAP as reverse micelles, to ensure the incorporation of DOTAP within the oily core.

**Figure 1:**
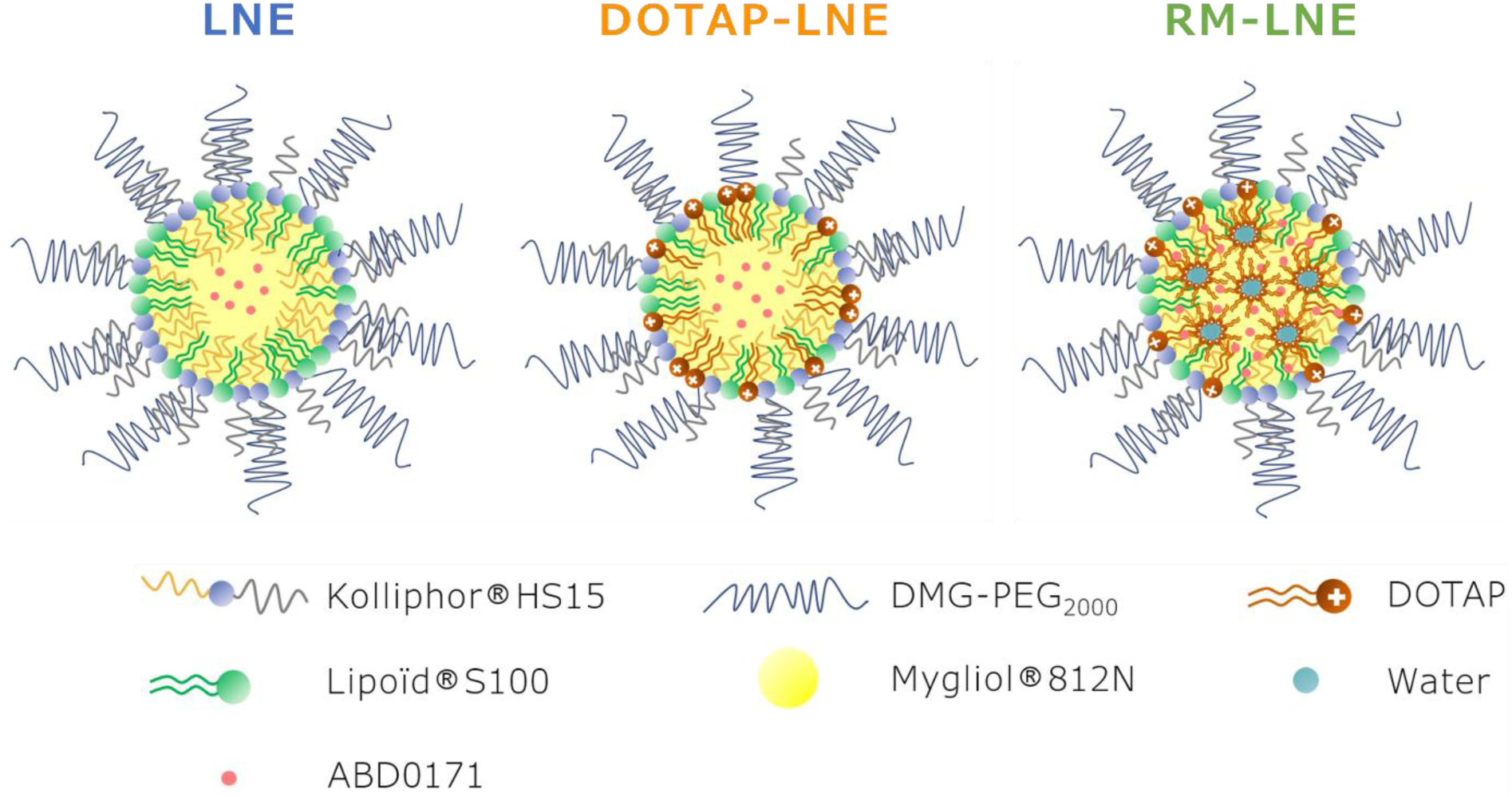
Schematic representation of LNE (blue), DOTAP-LNE (orange), and RM-LNE (green) formulations: LNE (without DOTAP), DOTAP-LNE (DOTAP directly mixed with other components in a one-pot process), and RM-LNE (DOTAP encapsulated as reverse micelles in the oily core).

This study demonstrates that lipid nanoemulsions incorporating the DOTAP cationic lipid are efficient and scalable nanocarriers for pulmonary drug delivery, showing robust physicochemical properties, high therapeutic efficacy against the H358 NSCLC cell line (IC50 < 5 µM), and significant tumor reduction *in ovo* while maintaining a favorable safety profile.

## EXPERIMENTAL SECTION

### Materials and reagents

The ALDH inhibitor (ALDHin) ABD0171 (S-methyl-4-[{2-[(3,4-dimethylbenzyl)oxy)ethyl}(methyl)amino]-4-methylpent-2-ynethioate) was synthesized and characterized by Roowin SAS (Riom, France) according to patent W02017064247. Kolliphor® HS15 (Macrogol (15)-hydroxystearate) was purchased from BASF (Ludwigshafen, Germany), Miglyol® 812N (triglycerides medium-chain) from IOI Oleochemical (Hambourg, Germany), Lipoïd® S100 (soy lecithin with 100% phosphatidylcholine) and DOTAP (1,2-dioleoyl-3-trimethylammonium-propane chloride salt) were purchased from Lipoid GmbH (Ludwigshafen, Germany). CF (5(6)-Carboxyfluorescein) was purchased from Merck (Darmstadt, Germany). DMPE-PEG_2000_ (N-(methylpolyoxyethylene oxycarbonyl)-1,2-dimyristoyl-sn-glycero-3-phosphoethanolamine sodium salt) was purchased from NOF EUROPE GmbH (Frankfurt, Germany). Deionized water was obtained using a Milli-Q Plus system (Millipore, Paris, France). All the reagents and solvents used for chromatography were of analytical grade. Cell culture media, fetal bovine serum (FBS), and buffers were purchased from VWR (Rosny sous bois, France). The NSCLC cell line, H358, was provided by the Institute of Advanced Biotechnologies (IAB), Grenoble. SYTOX Green and resazurin were purchased from Thermo Fisher Scientific (Massachusetts, USA).

### Preparation of Lipid NanoEmulsions (LNEs)

The detailed compositions of all the formulations are provided in the **Supporting Information (Table S1).**

The formulation of the lipid nanoemulsions (LNE) was adapted from the Phase Inversion Temperature (PIT) method reported by Garcion et al. (19). Briefly, the aqueous phase components (Kolliphor® HS15, Lipoïd® S100, and 0.9% NaCl) were mixed and added to the oil phase containing Miglyol® 812N, DMPE-PEG_2000_ and the ALDH inhibitor, ABD0171. The mixture was heated to 90°C and then cooled to 60°C, during which the phase inversion temperature (PIT) was observed. This heating-cooling cycle was repeated two additional times to ensure system stabilization. At the end of the final cycle, the formulation was cooled to the PIT (around 70°C), and a large volume of cold water was added to stabilize the emulsion, a step referred as "quenching". The formulation was stirred for 10 min at room temperature and stored at 2–8°C until use.

Based on this fabrication process, three distinct LNE formulations and their associated preparation methods were developed and optimized. Formulations were prepared with ABD0171 with a fixed drug loading of 2% (w/w) relative to the total lipid content, yielding a final drug concentration of 16 mg/mL. Formulations without drug were prepared following similar protocols and were referred as “vehicles”.

### Standard LNE formulation (LNE)

The standard LNE formulation corresponded to the PEGylated nanoemulsion described above encapsulating ABD0171.

### DOTAP LNE formulation (DOTAP-LNE)

The DOTAP-LNE formulation was prepared using the same process as described for the standard LNE formulation, except that DOTAP was added to the oil phase before the addition of the aqueous phase, to obtain a DOTAP-incorporating nanoemulsion.

### DOTAP reverse micelle LNE formulation (RM-LNE)

To prepare DOTAP reverse micelles (RM), Miglyol® 812N was mixed in a flask with DOTAP (previously dissolved in ethanol at 50 mg/mL) at a mass ratio of 5:1 (Miglyol® 812N:DOTAP). The mixture was placed in a rotary evaporator for 3 hours at 100 rpm, with a bath temperature of 50°C and a pressure of 95 mbar, to allow DOTAP to dissolve fully in Miglyol® 812N and ethanol evaporation. Next, water was added at a volume equivalent to 2% of the Miglyol® 812N-DOTAP mass, and the system was further rotated at 50°C and 100 rpm for 2 hours, enabling the organization of the reverse micelles. The reverse micelles were deemed ready once the mixture became fully translucent.

For RM-LNE fabrication, the LNE formulation excipients (Miglyol® 812N, Kolliphor® HS15, and Lipoid® S100) and the active molecule ABD0171 were mixed as previously described, followed by the thermocycling process. During the final heating cycle, when the system reached the last water in oil state at approximately 85°C, 735 mg of previously prepared DOTAP RM mixture were added. The system was then cooled down, and once the new phase transition temperature (PIT) of the system was observed (around 60°C), cold water was added (quenching) to halt the reaction and stabilize the system. The formulation was stored at 4°C before use.

### Evaluation of reverse micelle formation using 5(6)-carboxyfluorescein encapsulation

To evidence the formation of DOTAP reverse micelles (RM), 5(6)-carboxyfluorescein (CF) was encapsulated within the aqueous cores of ABD0171-loaded RM-LNE. RM(CF) formulations (with CF encapsulated in the aqueous droplets of the reverse micelles) were prepared by first dissolving CF in water at 50 mM, using 2–3 µL of 0.1 M NaOH to facilitate solubilization. This CF solution was added to the system after the 3 hours DOTAP solubilization in Miglyol® 812N, in the same proportions as for standard RM preparations (i.e., 2% by mass relative to the total Miglyol®-DOTAP mixture). ABD0171-loaded RM-LNE were then prepared as previously described. As a control, the same amount of CF was added at the final stage of DOTAP-LNE preparation (before the quenching step). For comparison with the free CF, the CF solution at 50 mM was first diluted in Milli-Q water to match the concentration used in the LNE formulations. This aqueous solution was then either measured directly (to compare with bulk formulations) or further diluted 1:2 (v/v) in ethanol (to mimic the disruption protocol applied to the LNEs). In both cases, final CF concentrations and dilution conditions were matched to those used in the corresponding formulation samples and served as references for maximal fluorescence intensity.

CF fluorescence measurements were conducted using a SpectraMax i3 plate reader (Molecular Devices, San Jose, CA, USA) with excitation at 450 nm, and emission spectra recorded from 475 to 800 nm in 2 nm increments. The samples (200 µL) were measured in black 96-well plates. To assess CF encapsulation, LNE formulations were disrupted by dilution in ethanol (1:2 v/v), followed by fluorescence measurement under the same conditions as the non-disrupted formulations diluted 1:2 v/v in MilliQ water.

### Characterization of LNE physicochemical properties

#### Measurement of size distribution and zeta potential by Zetasizer Nano ZS

The size distribution and zeta potential were measured using a Zetasizer Nano ZS instrument (Malvern Instruments, UK). Each formulation was diluted 1/100^th^ in milliQ water before measurement. The material refractive index was set to 1.449, material absorption to 0, dispersant viscosity to 0.8872 cP, dispersant refractive index to 1.33, and dispersant dielectric constant to 78.5. Measurements were performed after a 100 s equilibration period at 25°C using a 173° backscatter angle. Particle size measurements were conducted using plastic cuvettes, with three measurements per sample, each consisting of 10 runs of 10 seconds. Results were expressed as z-average particle diameter and polydispersity index (PDI), based on intensity distribution. Zeta potential was analyzed under the same dilution conditions using DTS1070 measurement cells. Three measurements per sample were performed, each comprising 20 runs. The dielectric constant of water was fixed at 78.5. Data were analyzed using Zetasizer Nano software and reported as mean values.

#### Cryo-Transmission Electron Microscopy (cryo-TEM)

3.5 μL of each sample were applied to glow-discharged Quantifoil 300 mesh R1.2/1.3 or R2/2 grids in a Vitrobot Mark IV (Thermo Fisher Scientific) at 22 °C and 100% humidity. After 10 seconds of incubation, excess liquid was blotted from both sides for 2–5 seconds using a blot force of −7 or 0, and grids were plunge-frozen in liquid ethane.

Grids were screened on a Glacios Cryo-TEM (Thermo Fisher Scientific) at the Institut de Biologie Structurale (IBS, Grenoble), operated at 200 kV. Regions containing thin vitreous ice and well-distributed particles were selected for data collection. Cryo-EM datasets were acquired using the EPU automated acquisition software. Micrographs were recorded using a Falcon 4i direct electron detector at nominal magnifications of 73,000×, 150,000×, or 310,000×, corresponding to calibrated pixel sizes of 1.9 Å, 0.94 Å, and 0.46 Å, respectively. All micrographs were acquired with a total dose of approximately 34 e⁻/Å² and defocus values between −2 and −3 μm. Images were analyzed using ImageJ for nanoparticle size measurements, as described in supplementary information **Figure S3**.

#### Quantification of ABD0171

Quantification of ABD0171 was performed using an Ultimate 3000 High-Performance Liquid Chromatography (HPLC) chain from Thermo Fisher. The extraction of ABD0171 from the formulations was performed by dissolving 140 µL of sample with 22.85 mL acetonitrile, achieving a final dilution of 1:164. The resulting solution was transparent, and 1 mL was transferred to HPLC vial for analysis. Analysis was performed using a Waters Symmetry C8 analytical column (4.6 × 50 mm, 3.5 µm) maintained at 40 °C, with a modified mobile phase composition. The mobile phase consisted of 10 mM ammonium acetate (phase A) and acetonitrile (phase B) in an isocratic ratio of 50:50 (v/v). The flow rate was set to 1.5 mL/min and UV detection was performed at 270 nm. The injection volume was 5 µL with a total run time of 10 min. The retention time for ABD0171 was 7.74 ± 1.0 min. External calibration was performed based on analytical standards. A calibration curve was established and demonstrated a linear response (R² > 0.99) across the tested concentration range (3.6 - 644 µg/mL). Data analysis was conducted using the Chromeleon software (Thermo Fisher, Waltham, MA, USA).

#### Characterization of LNEs formulations by Taylor Dispersion Analysis

SD-TDA (Size Distribution - Taylor Dispersion Analysis) experiments were performed on a TaylorSizer instrument from Nanoscale Metrix (Toulouse, France) equipped with a diode array detector set at 200 nm and 270 nm, for the detection of RM and ABD0171, respectively. Bare-fused silica capillaries with inner diameter of 50 μm, a total length of 60 cm, and a detection window length of 49.5 cm, were used for all experiments. The temperature of the capillary area was set to 25°C for the LNE formulations and 35°C for RM sample. The data processing was performed by TaylorSoft which included a patented algorithm enabling size distribution through SD-TDA.

#### *In vitro* release of ABD0171 from LNEs formulations

The release of ABD0171 from the formulations was assessed by dialysis using Slide-A-Lyzer^TM^ Dialysis Cassettes (Thermo Fisher, 10 kDa MWCO, 3 mL capacity). Following hydration of the membrane, 3 mL of the formulation were loaded into the dialysis cassettes and immersed in 400 mL of 0.9% NaCl solution (release medium) under gentle stirring to maintain homogeneous conditions. The large volume of the release medium relative to the sample ensured the sink conditions throughout the experiment. After overnight incubation, the concentration of the retained drug in the dialysis cassette was quantified using previously described high-performance liquid chromatography (HPLC) method.

#### Stability study of LNE formulations

The stability of the formulations was studied at 37°C, room temperature (RT), and 4°C. Each formulation was stored in glass vials, and at predetermined time points, the stability of the formulation was evaluated by measuring the physicochemical characteristics by DLS (colloidal stability) and by HPLC (ABD0171 quantification).

### *In vitro* evaluation

#### Cell line and culture conditions

H358 cells were cultured in RPMI 1640 medium supplemented with 1% L-glutamine, 1% penicillin/streptomycin, and 10% FBS unless otherwise specified. Cells were maintained in a humidified atmosphere containing 5% CO_2_ at 37°C.

#### Cytotoxicity assessment

The growth inhibitory effects of the formulations were evaluated using a resazurin-based cytotoxicity assay. H358 cells were seeded into 96-well plates at a density of 22,000 cells/cm^2^ (7,000 cells/well) and incubated overnight at 37°C with 5% CO₂. For each evaluated formulation, cells were treated with a range of ABD0171 concentrations (1 – 32 µM) and incubated at 37°C with 5% CO_2_ for 48 h. Untreated cells (cell culture medium only) constituted the 100% viability control. A 20 µL volume of a 1X resazurin solution, prepared by diluting a 10X stock solution (1.5 mg/mL) in PBS, was added to each well and incubated for 3 h at 37°C. The fluorescence was measured (excitation: 560 nm; emission: 590 nm) using a SpectraMax i3 plate reader. The half-maximal inhibitory concentration (IC₅₀) was determined via non-linear regression analysis of log-dose/response curves using the GraphPad^TM^ software. For each formulation, the experiment was repeated with N = 3 different formulations, with five biological replicates each time.

#### Cell viability assessment by flow cytometry (FACS)

Cell viability was assessed by flow cytometry based on membrane integrity loss using the SYTOX^TM^ Green marker, which only penetrates dead cells with compromised membranes. H358 cells were seeded in 24-well plates at a density of 21,000 cells/cm^2^ (40,000 cells/well) and incubated overnight at 37°C with 5% CO2. For each evaluated formulation, cells were treated with ABD0171 concentrations of 1.5 μM and 6 μM for 2, 4, 6, and 24 h. At each time point, the medium was collected, and the cells were washed, detached with trypsin, and centrifuged at 180 × g for 5 min. The cell pellet was resuspended in PBS and stained with 5 μM SYTOX Green for 10 min in the dark. The fluorescence intensity was measured using a BD Accuri C6 Plus flow cytometer (Becton Dickinson) with excitation at 488 nm and emission at 585/40 nm (FL2 filter). The data were analyzed using the Kaluza^TM^ software and expressed as the percentage of dead cells. For each formulation, the experiment was repeated with N = 3 different formulations.

### *In vivo* evaluation

#### Efficacy and toxicity study in chicken ChorioAllantoic Membrane (CAM) model

According to French legislation, no ethical approval is required for experiments using oviparous embryos (Decree No. 2013–118, February 1, 2013; Article R-214-88). The chicken CAM assay was performed by Inovotion SAS (La Tronche, France) using fertilized White Leghorn eggs (Couvoir Hubert, France). The eggs were incubated at 37.5°C with 50% relative humidity for 9 days. On EDD9 (Embryonic Development Day 9), the chorioallantoic membrane (CAM) was dropped down by drilling a small hole into the air sac, followed by creating a 1 cm² window in the eggshell above the CAM. At least 15 embryos per group were used in the experiments.

The H358 tumor cell line was cultured in RPMI 1640 medium supplemented with 10% FBS, 1% sodium pyruvate, and 1% penicillin/streptomycin. On EDD9, H358 cells were detached using trypsin, washed in complete medium, and resuspended in graft medium. A total inoculum of 1 × 10⁶ cells was applied onto the CAM of each egg, after which the eggs were returned to the incubator. Eggs were randomly divided into five treatment groups: negative control (0.9% NaCl), ABD0171 free compound, LNE, DOTAP-LNE, and RM-LNE. Treatments were administered at two ABD0171 doses of 2.76 mg/kg [1X] and 27.6 mg/kg [10X] by dropwise application onto the tumors on EDD11 and EDD13. Egg survival was monitored daily until the end of the experiment. On EDD18, embryos were sacrificed, and the upper portion of the CAM containing the tumor was excised, washed with PBS, and fixed in 4% paraformaldehyde (PFA) for 48 hours. Tumors were carefully separated from the surrounding CAM tissue and weighed to assess the average tumor weight in the experimental groups.

#### Toxicity study in mice

The toxicity studies were conducted in collaboration with the Optimal Platform (La Tronche, France). Mice experiments were conducted in compliance with protocols approved by the Grenoble Ethics Committee and the French Ministry of Higher Education and Research (APAFIS#33137-2021110411585349v2). Forty-three six-week-old female NMRI nude mice (Janvier Labs, France) were used in this study. The details of the experimental groups and treatments are presented in **Figure S1**.

Formulations were prepared by dilution in 0.9% NaCl and administered via tail vein intravenous bolus injections (8 µL/g). To determine the maximum tolerated dose, body weight, behavior, and distress were monitored with euthanasia performed if ethical limits were exceeded. All the mice were euthanized for organ analysis at the end of the study.

### Statistical Analysis

All statistical analyses were conducted using the GraphPad^TM^ Prism software (version 10.5.0). Data are presented as mean ± standard deviation (SD). For *in vitro* experiments, statistical comparisons between groups were performed using one-way ANOVA. Results were considered statistically significant at p < 0.05. For the chicken CAM assay, efficacy was analyzed using one-way ANOVA with Tukey’s multiple comparisons test, and survival was assessed using the Log-rank (Mantel-Cox) test. Statistical differences between groups are indicated on graphs using a star notation system: *0.05 ≥ p > 0.01, **0.01 ≥ p > 0.001, *** 0.001 ≥ p > 0.0001 and **** p ≤ 0.0001.

## RESULTS AND DISCUSSION

To enable the efficient delivery of the lipophilic ALDH inhibitor ABD0171 we developed and optimized lipid nanoemulsions (LNEs) capable of stably incorporating this compound into their oily core. LNEs represent a promising class of nanocarriers due to their biocompatibility, high drug-loading capacity, and scalable manufacturing processes. In this study, we systematically explored formulation parameters to generate monodisperse and stable nanoemulsions suitable for pulmonary administration of ABD0171. In parallel, we investigated the incorporation of the cationic lipid DOTAP as a targeting element to promote lung tropism. Three distinct LNE formulations were designed: a simplified baseline LNE, a DOTAP-enriched LNE (DOTAP-LNE), and a DOTAP reverse micelle–based LNE (RM-LNE). Particular attention was given to the incorporation of the cationic lipid DOTAP, either directly at the oil–water interface or via preformed reverse micelles (RMs), to modulate surface charge and drug encapsulation behavior. These two incorporation strategies were designed to address distinct hypotheses. The RM-LNE aimed to localize DOTAP within the oily core, mimicking the internal structure of LNPs used for mRNA delivery, for which core-localized DOTAP have been shown to be critical for efficient lung organ targeting (15). In contrast, the DOTAP-LNE allowed DOTAP to remain exposed at the surface, providing a simpler alternative that could still enable surface-mediated interactions relevant for pulmonary targeting. The simplified LNE served as a reference to evaluate the impact of DOTAP incorporation strategies on colloidal properties and delivery efficiency. The hypothesized structures of these formulations, including the localization of DOTAP, are illustrated in **Figure 1** along with the commercial names of their components. The detailed mass proportion of each formulation is presented in **Supplementary Information, Table S1.**

### Development and fluorescent confirmation of DOTAP reverse-micelles lipid nanoemulsions (RM-LNE)

The RM-LNE formulation was developed based on the solvent-free thermocycling method proposed by Heurtault et al. (18). The selected composition of the formulation employs FDA-approved ingredients for parenteral administration, ensuring the safety and clinical translatability of the resulting formulation. To enhance pulmonary delivery, DOTAP was incorporated into the core of the nanoemulsion according to the results outlined by Dilliard et al. (15). In contrast to Dilliard’s microfluidic manufacturing strategy, we employed thermocycling, which requires an innovative approach to ensure the effective integration of DOTAP into the oily core of the nanoemulsion. Drawing from the work of Vrignaud et al., who demonstrated the incorporation of anionic dioctylsodium sulfosuccinate (AOT) micelles into lipid nanocapsules for peptide delivery (20–22), we investigated the formation of cationic reverse micelles to incorporate DOTAP. Jörgensen et al. demonstrated that reverse micelles formed with didodecyldimethylammonium bromide offered significant advantages over AOT-based formulations, including improved encapsulation efficiency, greater stability, enhanced bioavailability, and stronger membrane interactions (23).

The key steps in the manufacturing process of the RM-LNE formulation are illustrated in **Figure 2**, highlighting the optimization process parameters evaluated during formulation development. These parameters included the lipid concentration of the nanoemulsion, the amount of DOTAP in the reverse micelle structure, the water-to-lipid ratio in the reverse micelle structure, and the volume of reverse micelles added to the lipid nanoemulsion. The optimized process parameters included a lipid concentration of 252 mg/mL, a DOTAP:Miglyol® 812N weight ratio of 5:1, 2% (w/w) of water in the RM formulation, and 735 mg of RMs were added per 5 mL of total formulation.

**Figure 2:**
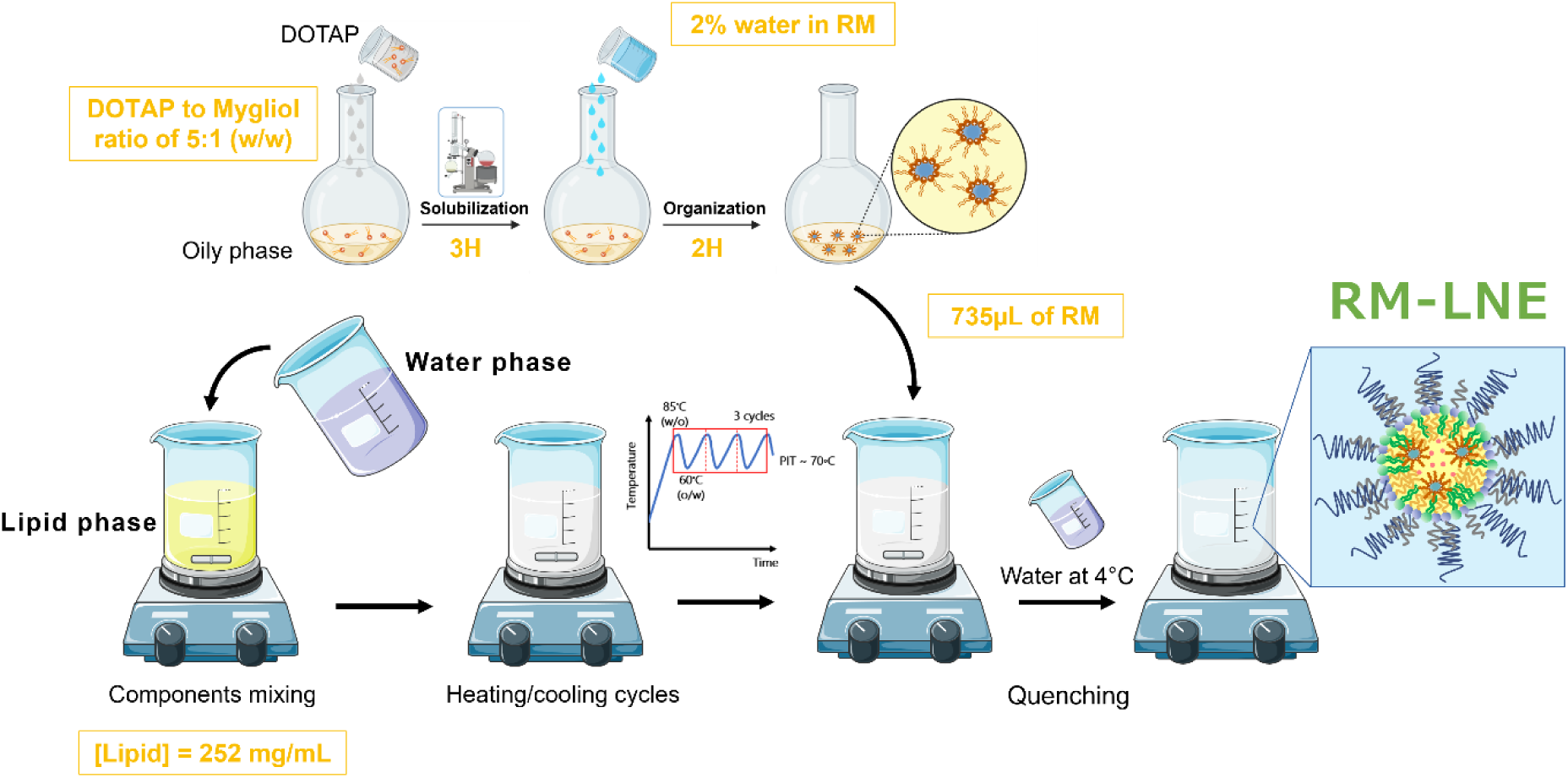
Scheme illustrating the key steps in the preparation of DOTAP reverse-micelle lipid nanoemulsions (RM-LNE), with the optimized process parameters highlighted in orange. The optimization particularly focused on the DOTAP content in the RM structure, the water percentage within the RM, and the volume of RM added to the LNE formulation. RM-LNE formulations were fabricated using a thermocycling process, with reverse micelles (RMs) incorporated during the final temperature cycle.

The formation of RM was verified by DLS, which provided insights into their size distribution, polydispersity, and stability. The data confirmed the formation of RMs with a monodisperse size distribution and a mean diameter of 14 nm (**Supplementary Information Figure S2**), which was consistent with previous reports on RM characterization (24).

To investigate the effective incorporation of RM into LNEs, we used 5(6)-carboxyfluorescein (CF), a hydrophilic dye commonly used as a fluorescent tracer in nanoparticle systems. CF is widely recognized for its ability to self-quench at high concentrations, due to dimerization and energy transfer mechanisms, making it a reliable marker for encapsulation studies (25). CF was either incorporated into the aqueous core of RM prior to LNE formulation (RM(CF)-LNE) or added at the end of the thermocycling process in a formulation without RM structure (DOTAP(CF)-LNE). In this control, CF was expected to reside in the aqueous dispersant phase and not in the oily core of the LNE. A solution of CF, prepared at the same concentration and subjected to the same dilution conditions (in water or ethanol), was used as a reference for maximal fluorescence (**Figure 3A)**.

**Figure 3:**
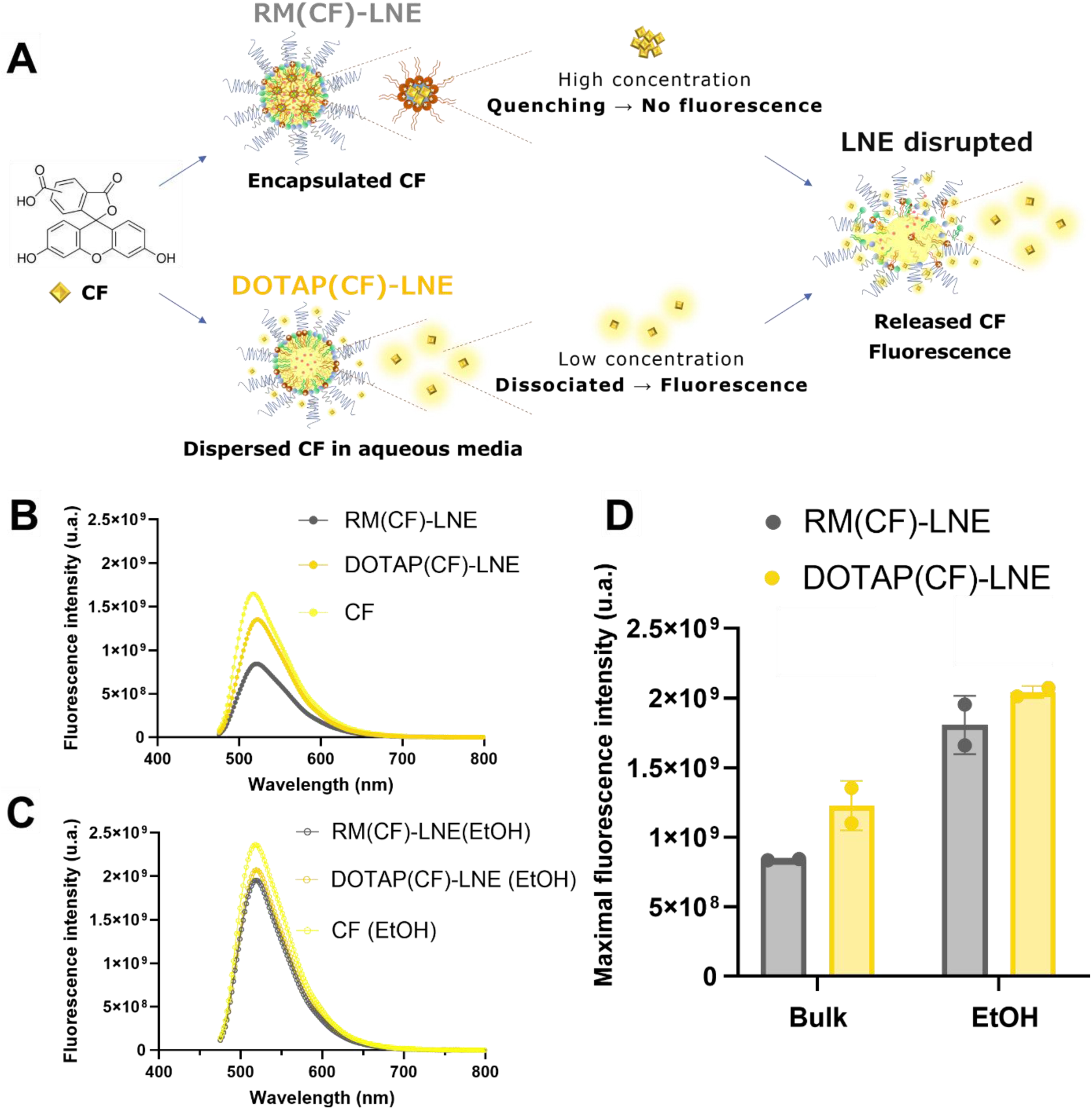
Reverse micelle encapsulation assay using CF and fluorescence analysis. **(A)** Development of a reverse micelle encapsulation assay using 5(6)-carboxyfluorescein (CF). RM(CF)-LNE constitutes the formulation with DOTAP reverse micelles encapsulating CF, whereas DOTAP(CF)-LNE includes DOTAP and CF directly added to the other components without the RM structure. **(B-C)** Fluorescence emission spectra of both formulations compared to free CF at equivalent concentrations. Spectra were recorded in bulk aqueous medium (B) or after ethanol disruption (C). Free CF in water (B) or in ethanol (C) is included as a reference. **(D)** Maximal fluorescence intensity (Em ∼ 515 nm) of RM(CF)-LNE and DOTAP(CF)-LNE formulations in bulk condition and after disruption in ethanol.

The size distribution of RM and RM(CF), provided in **Supplementary Information Figure S2**, demonstrated that the CF incorporation into the RM structure did not affect its size distribution. As shown in **Figure 3B**, the RM(CF)-LNE formulation displayed a markedly lower fluorescence spectra compared to both the DOTAP(CF)-LNE control and the free CF reference, consistent with self-quenching of CF molecules concentrated within the aqueous cores of reverse micelles. In contrast, the fluorescence intensity of DOTAP(CF)-LNE was closer to that of free CF, suggesting that CF in this control formulation was predominantly dispersed in the aqueous phase, where it remained largely unquenched. This finding was consistent with previous studies showing that reverse micelles can effectively encapsulate hydrophilic molecules, enabling controlled release and enhanced stability (17). After ethanol disruption of both RM(CF)-LNE and DOTAP(CF)-LNE formulations (**Figure 3C**), the fluorescence spectra of RM(CF)-LNE increased substantially and became comparable to that of the DOTAP(CF)-LNE and free CF, indicating the release and dilution of CF from the micellar cores into the aqueous phase. Notably, fluorescence values in ethanol were consistently higher than in water, including for the free CF reference, likely due to improved solubilization and reduced self-quenching in the organic solvent. This was confirmed quantitatively in **Figure 3D**, where the maximal fluorescence of RM(CF)-LNE was lower than that of DOTAP(CF)-LNE in the bulk condition, but remained nearly identical after ethanol disruption. Additionally, both formulations showed significantly higher fluorescence after disruption compared to their bulk counterparts, further supporting the combined effects of nanoparticle dissociation and enhanced CF solubilization in ethanol. These findings validated the successful incorporation of CF into the oily core of LNEs via DOTAP reverse micelles. This approach not only highlights the robustness of RM formation and their integration into LNEs but also underscores the potential of this platform for advanced drug delivery applications.

### Physicochemical characterization of LNEs

The physico-chemical properties of the three different LNE vehicles (LNE, DOTAP-LNE, RM-LNE) loaded with ABD0171 were characterized by DLS and cryo-TEM.

During formulation optimization, we evaluated the effect of increasing DOTAP content on particle size in both one-pot and RM methods. Regardless of the strategy, increasing the proportion of DOTAP induced a gradual increase in diameter (**Supporting Information, Figure S4A**). These experiments were performed with constant drug loading (Miglyol®: ABD0171 mass ratio 5:1), suggesting that the observed increase in size was driven by formulation parameters rather than drug content. The size increase is likely related to the localization of DOTAP at the nanoparticle surface, a position that should favor its interaction with Kolliphor® HS15, causing a more pronounced impact on particle size (26,27). In particular, the larger polar head of DOTAP increases the effective head group area (a₀), reducing the packaging parameter (P), and consequently increasing the curvature radius and thus the particle size (28,29). Based on these findings, we selected optimized formulation parameter for further comparison (**Table S1**).

Size measurement by DLS **(Figure 4A)** revealed that the RM-LNE formulation as a vehicle (without encapsulated ABD0171) demonstrated a higher diameter (60 ± 10 nm), though not statistically significant, compared to its counterpart LNE (50 ± 0.7 nm) and DOTAP-LNE (49 ± 0.7 nm). Incorporating ABD0171 significantly increased the particle size in the DOTAP-LNE formulation from 49 nm to 72 nm, whereas drug encapsulation did not induce any significant increase in diameter for LNE and RM-LNE. This observation may suggest that ABD0171 incorporation induces interfacial rearrangements, particularly in the DOTAP-LNE formulation. Lipophilic compounds can partition into the lipid core or interfacial region, disrupting lipid organization and altering the system’s hydrophilic-lipophilic balance (30). Such perturbations may lead surface-localized DOTAP to rearrange or interact more strongly with surfactants like Kolliphor® HS15, resulting in a thicker interfacial layer and increased droplet size. A similar mechanism has been proposed in studies showing that DOTAP incorporation into liposomes increases particle size and reduces lipid crystallinity by modulating membrane organization (31).

**Figure 4:**
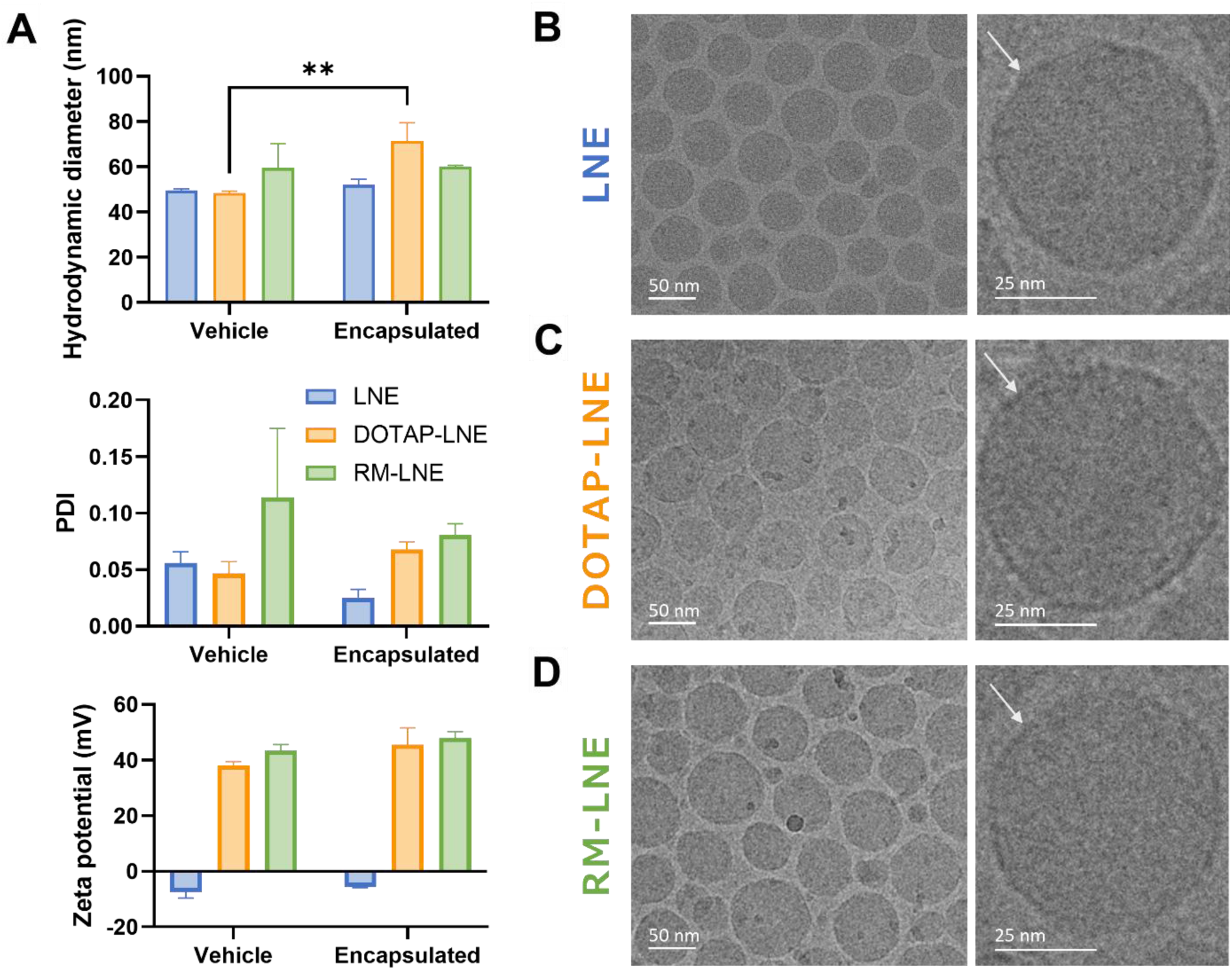
Physicochemical comparison of LNE, DOTAP-LNE, and RM-LNE formulations with and without ABD0171 encapsulation. (A) Physicochemical characteristics of formulations, including hydrodynamic diameter (nm), polydispersity index (PDI), and zeta potential (mV), as determined by dynamic light scattering (DLS). Data are expressed as the mean ± standard deviation (SD) from three independent measurements (n=3). (B-D) Representative cryo-TEM images of ABD0171-loaded formulations: (B) LNE, (C) DOTAP-LNE, and (D) RM-LNE. Left panels show overview images; right panels display high-resolution views of individual nanoemulsion droplets. White arrows indicate the lipid shell surrounding the nanocarriers.

In contrast, RM-LNE incorporates DOTAP partially within reverse micelles in the oil phase. This pre-organization may reduce the structural impact of drug loading. As a result, the interfacial layer of RM-LNE requires less readjustment, which may help maintain the integrity of the surfactant layer and thereby limiting size variation. To be noted that all formulations exhibited PDI values below 0.15, confirming monodispersity and demonstrating that drug encapsulation did not compromise size distribution or induce aggregation.

These size trends were also confirmed by cryo-TEM imaging (**Figure 4B–D**), which revealed well-defined, monodisperse populations of spherical nanoparticles for all formulations, in agreement with DLS data. The measured size distribution by cryo-TEM further confirmed the successful formation of a single-size population around 55 nm. To build on the cryo-TEM–derived size distributions, we implemented a first-pass, image-based estimation of particle concentration (particles/mL). We relied on manual nanoemulsion counting within a defined two-dimensional region of interest (ROI), followed by extrapolation to a three-dimensional volume under the assumption of uniform spherical geometry. The detailed workflow is outlined in **supporting information Figure S3**. This approach yielded estimated concentrations of 8.1 × 10¹⁴, 6.0 × 10¹⁴, and 7.3 × 10¹⁴ particles/mL for LNE, DOTAP-LNE, and RM-LNE, respectively, values consistent with their similar average diameters (∼ 55 nm). Notably, cryo-TEM has previously been shown to yield statistically robust nanoparticle distributions that closely match those obtained by standard particle-counting techniques. For instance, Nordström et al. (32) employed quantitative cryo-TEM to characterize PEGylated liposomal doxorubicin (Doxil®-like), reporting a strong correlation with Nanoparticle Tracking Analysis (NTA) and DLS derived values, thereby supporting the potential of cryo-TEM for reliable nanoparticle concentration quantification. No aggregates nor secondary populations were observed, supporting the low PDI values (< 0.15) obtained by DLS **(Figure 4A)**. Furthermore, close-up images (right panels) clearly show a distinct lipid shell surrounding the oily core (indicated by white arrows), highlighting the structural integrity of the nanoemulsion. No notable morphological differences were observed between the DOTAP-LNE and RM-LNE formulations, despite their differing methods of DOTAP incorporation (surface-localized vs. reverse micelle strategy) **(Figure 4C and D, respectively)**. In particular, no micelle-like structures could be visualized within the lipid core of RM-LNE, suggesting either their disassembly or integration into the matrix below the resolution threshold of conventional cryo-TEM (∼ 5 nm under our imaging conditions). To further investigate the internal organization of our nanoemulsions, advanced imaging techniques such as cryo-electron tomography (cryo-ET), known to reconstruct 3D nanoparticle architecture from tilt-series data (33), could provide definitive evidence of compartmentalization or hidden reverse micelles within the oily core.

Zeta potential (ZP) measurements **(Figure 4A)** indicated a stable positive surface charge (+45 mV) for both DOTAP-LNE and RM-LNE, regardless of DOTAP incorporation strategy. It is worth noting that zeta potential reflects the electric potential at the slipping plane rather than the actual surface charge density. As such, although increasing the amount of DOTAP at the interface effectively raises the number of positively charged headgroups, this does not necessarily result in a higher zeta potential. This may be due to electrostatic screening by counter-ions or to the fact that, beyond a certain threshold, the interfacial region cannot sustain a stronger electric field at the slipping plane. Supporting data from **Figure S4B** reinforces this hypothesis, as the increase in zeta potential with DOTAP content became progressively less pronounced beyond ∼6% (w/w) for both formulations. This trend suggests that, beyond a certain threshold, additional DOTAP has limited influence on the electrokinetic potential. These findings align with studies showing charge density saturation at the nanoparticle surface, where additional charged components did not further increase ZP (34,35).

In summary, the distinct positioning of DOTAP significantly affected the nanoparticle size, likely through its interactions with Kolliphor® HS15, while the observed charge behavior reflected a saturation effect. These results provide valuable insights into the structural and electrostatic factors influencing the design and stability of functional LNEs.

### Encapsulation yield and encapsulation efficiency of ABD0171 in lipid nanoemulsions (LNEs)

Encapsulation yield and encapsulation efficiency are critical parameters that influence the performance of lipid-based drug delivery systems. In this study, all nanoemulsion formulations were prepared with a fixed ABD0171 drug loading of 2% (w/w) relative to total lipid content, allowing direct comparison across the different LNE platforms. Encapsulation yield refers to the final quantity of drug in the formulation compared to the initial amount of drug, translating the efficacy of the fabrication process to avoid drug loss. The encapsulation yields of the optimized formulations (LNE, DOTAP-LNE, and RM-LNE) were assessed using HPLC.

All formulations encapsulating ABD0171 **(Figure 5A)** exhibited yields approaching 100% **(Figure 5B).** This high yield can be explained by the absence of a purification step after nanoemulsion formation, meaning that both encapsulated ABD0171 and any non-associated components, such as free drug, unincorporated surfactants, or small micellar species, remained in the final product. In preclinical studies, purification is commonly implemented to minimize the presence of unbound components that may alter biodistribution or raise toxicity concerns. While such steps can improve the safety profile of the final product, they also increase processing time and manufacturing costs and may induce significant drug loss. Notably, non-purified nanoemulsion may be more therapeutically relevant for clinical use, provided that the safety of all excipients and the overall composition is rigorously demonstrated. In line with this strategy, we chose not to perform a purification step in the present study, allowing for a realistic evaluation of the formulation in its entirety, including non-encapsulated species. In addition, the consistently high recovery of ABD0171 across formulations also suggests that the compound remained chemically stable throughout the thermocycling process. This observation is consistent with its high lipophilicity and likely preferential partitioning into the oily core, which would limit its exposure to hydrolytic degradation during processing.

**Figure 5:**
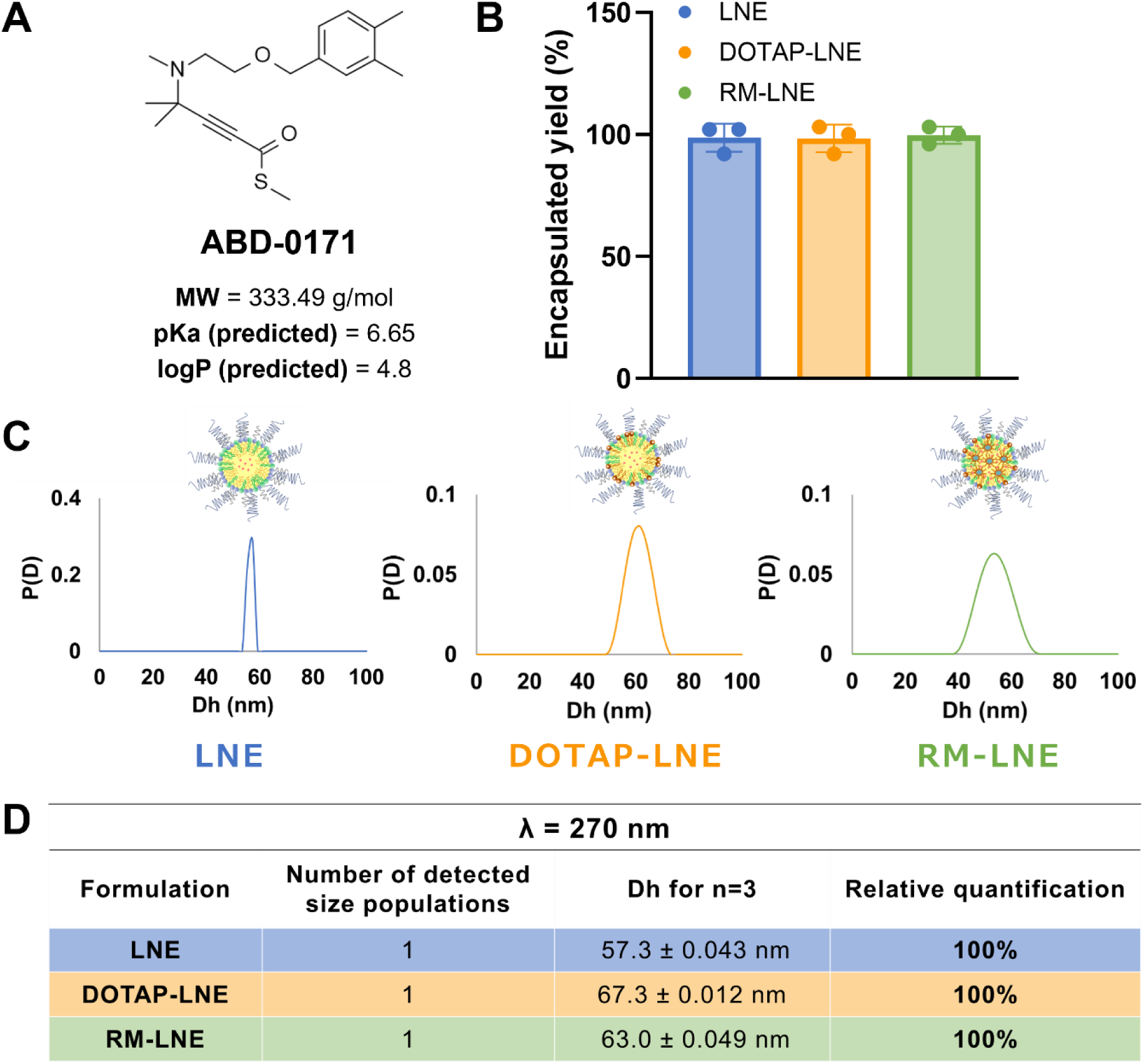
Encapsulation performances of ABD0171 in LNE formulations. **(A)** Chemical structures of ABD0171, highlighting key physicochemical properties, such as molecular weight, predicted pKa, and predicted log P, relevant to this formulation and biological activity. **(B)** Encapsulation yield (%) of ABD0171 in LNE (blue), DOTAP-LNE (orange), and RM-LNE (green) formulations (mean ± SD, n = 3). **(C)** SD-TDA spectral profiles of LNE, DOTAP-LNE and RM-LNE. **(D)** Population distribution (Dh, hydrodynamic diameter) and relative quantification of ABD0171 in the main population, for each formulation, as determined by SD-TDA.

To complement these results, we next quantified the actual encapsulation efficiency (EE%) to evaluate how effectively ABD0171 was encapsulated within the lipid core of the nanoemulsions. While recovery yield reflects the total amount of drug present in the final nanoemulsion, EE% specifically indicates the fraction truly incorporated within the nanostructure, rather than freely dispersed in the surrounding medium.

To assess the EE%, Size Distribution Taylor Dispersion Analysis (SD-TDA) was performed at 270 nm, the ABD0171 absorption wavelength. As shown in **Figure 5C** and **5D**, we observed that the drug is fully incorporated into the lipid core of the nanoemulsions for all types of LNEs. This high encapsulation efficiency is likely attributable to the compound physicochemical properties, particularly its high lipophilicity, as reflected by its predicted logP value (6.65). Roger et al. (36) previously highlighted that surfactants such as Kolliphor® HS15 can form residual micellar structures even after nanoemulsion formation. Their work further emphasized that these micelles may contribute to drug encapsulation, especially for compounds with higher affinity for aqueous or interfacial environments. However, this behavior did not appear to take place during the encapsulation of ABD0171, likely due to its strong lipid affinity, which favored partitioning into the oily core over the aqueous phase or the surfactant micelles.

To further confirm the structural integration of DOTAP RM within the lipid core, we performed SD-TDA at 200 nm, a wavelength specific to RM absorption. At this wavelength, free RM dispersed in Miglyol® 812N displayed a narrow distribution centered around 14 nm (**Supporting Information Figure S5A**), in line with DLS data (**Supporting Information Figure S2**). In contrast, the RM-LNE formulation showed a single peak at ∼60 nm (**Figure S5B**), indicating that the RM specific absorbance was now associated with the nanoemulsion droplets. This diameter shift, combined with the presence of a single population, provides compelling evidence that RM structures are not freely dispersed in solution, but indeed embedded within the lipid core. These results are in accordance with the results obtained at 270 nm for the ABD0171 profile (**Figure 5D**) and together validate the efficient co-localization of both DOTAP RM and the drug within the same nanostructured carrier.

### *In vitro* release and long-term stability of ABD0171 in nanoemulsions

The drug release profiles of the LNE formulations were evaluated by dialysis for 24 h. As shown in **Figure 6A**, ABD0171 was largely retained in the lipid core of all nanoemulsions, with minimal release observed after 24 h. The high lipid affinity of ABD0171 made the drug preferentially associated with the lipid core of the nanoemulsions, leading to better retention and slower release. This was in accordance with the SD-TDA results (**Figure 5B** and **C**), which showed that 100% of ABD0171 was effectively encapsulated in the oily core of the nanoemulsion. These findings were consistent with those of previous studies demonstrating how the chemical properties of drugs influence their distribution within lipid nanoparticles and their release profile. For example, Knoke et al. observed that lipophilic drugs are preferentially localized within the lipid core of nanoemulsions, where they are more stably retained, whereas hydrophilic drugs are more likely to migrate toward the aqueous phase, resulting in faster release profiles (37). Similarly, the structural insights provided by Spinozzi et al. further confirmed that the internal organization of lipid nanoparticles is directly influenced by the lipophilicity of the incorporated compounds, which dictates their localization and retention (38).

**Figure 6:**
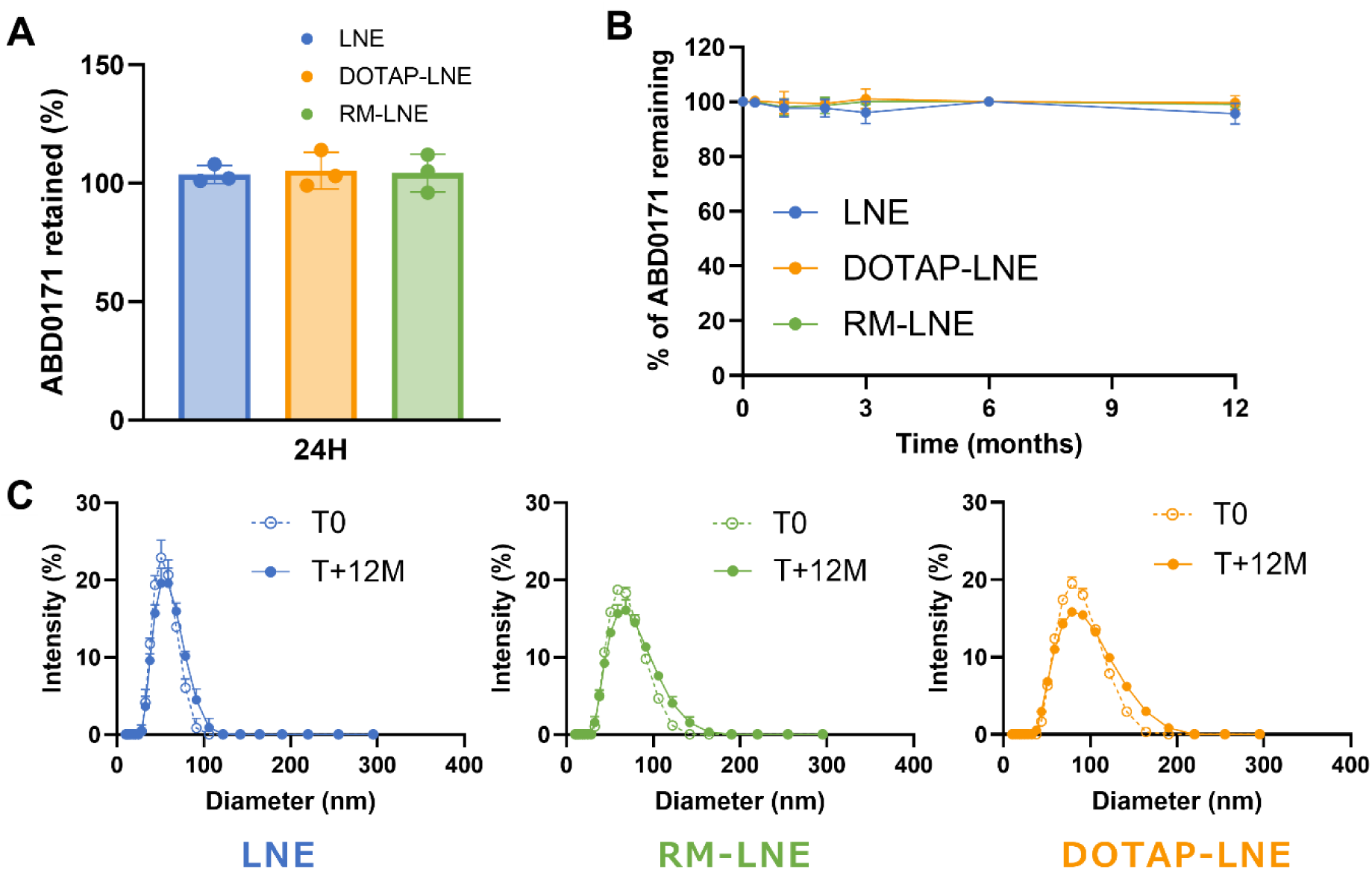
Stability assessment of ABD0171 in LNE formulations: retention, chemical, and colloidal stability. **(A)** Retention percentages of ABD0171 after 24 h of dialysis. **(B)** Chemical stability of ABD0171 at 4°C for a 12 month period, as quantified by HPLC. **(C)** Colloidal stability of nanoemulsions after 12 months of storage at 4°C, assessed by size distribution measurements using DLS.

The stability of the formulations was further evaluated after 12 months of storage at 4°C. As shown in **Figure 6B**, the chemical stability of ABD0171 was preserved across all formulations, with only minor losses (< 5%) observed after 12 months. Dynamic Light Scattering (DLS) analysis (**Figure 6C**) confirmed the colloidal stability of all formulations, with no evolution in size distribution and a PDI consistently below 0.1, indicating that all nanoemulsions remained stable during the storage period. These results aligned with those of Danaei et al.,(39) who emphasized the critical role of particle size and polydispersity index (PDI) in determining the stability of lipid-based nanoparticles. A low PDI, as observed in our formulations, is essential for maintaining colloidal stability, ensuring uniform particle size, and minimizing aggregation and Ostwald ripening during prolonged storage (39). The consistent colloidal properties observed in our study further reinforced the robustness of the formulations under storage conditions, even for storage periods as long as 12 months, which is necessary for clinical translation.

### *In vitro* evaluation of therapeutic efficacy and safety of ABD0171 LNE formulations

To demonstrate the therapeutic potential of our approach for lung cancer treatment, we first assessed the *in vitro* efficacy of ABD0171 encapsulated in lipid nanoemulsions (LNE, DOTAP-LNE, and RM-LNE) in H358 NSCLC cells using a resazurin assay. The IC50 values (**Figure 7A** and **7B**) for all formulations were consistently below 5 µM, regardless of the zeta potential or composition of the formulation. These results suggested strong therapeutic potential, as IC50 values below 5 µM are typically indicative of highly promising anticancer agents (40,41). IC50 values of the formulations were slightly lower compared to the free molecule, although not statistically different, which is commonly observed in *in vitro* settings for which formulation benefits (e.g., improved pharmacokinetics and distribution profiles) cannot be fully exploited. To further evaluate the efficacy of the formulations, we assessed the cell death profiles using flow cytometry and SYTOX staining. **Figure 7C** displays the histograms of SYTOX-positive (SYTOX+) cell populations, which provides the quantification of dead cells expressed as a percentage of the total population. Additionally, the kinetics of cell death overtime were monitored and are presented in **Figure 7D**, where the percentage of dead cells is plotted as a function of treatment concentration and time. Notably, a marked increase in cell death was observed over time at a concentration of 6 µM compared to 1.5 µM, for which the dead cell population remained stable and low (< 10%) throughout the observation period, in accordance with the IC50 value of ∼ 5 µM. These findings confirmed the concentration-dependent nature of the cytotoxic effects, likely reflecting a progressive inhibition of ALDH activity with increasing drug levels. In addition, the time-dependent increase in cell death at concentrations above the IC_50_ suggests a delayed cytotoxic effect consistent with the mechanism of action of ABD0171: as an ALDH inhibitor, ABD0171 efficacy may rely on the gradual accumulation of toxic aldehyde species ultimately inducing apoptosis, leading to cell death over time. These trends were consistent across all tested formulations.

**Figure 7:**
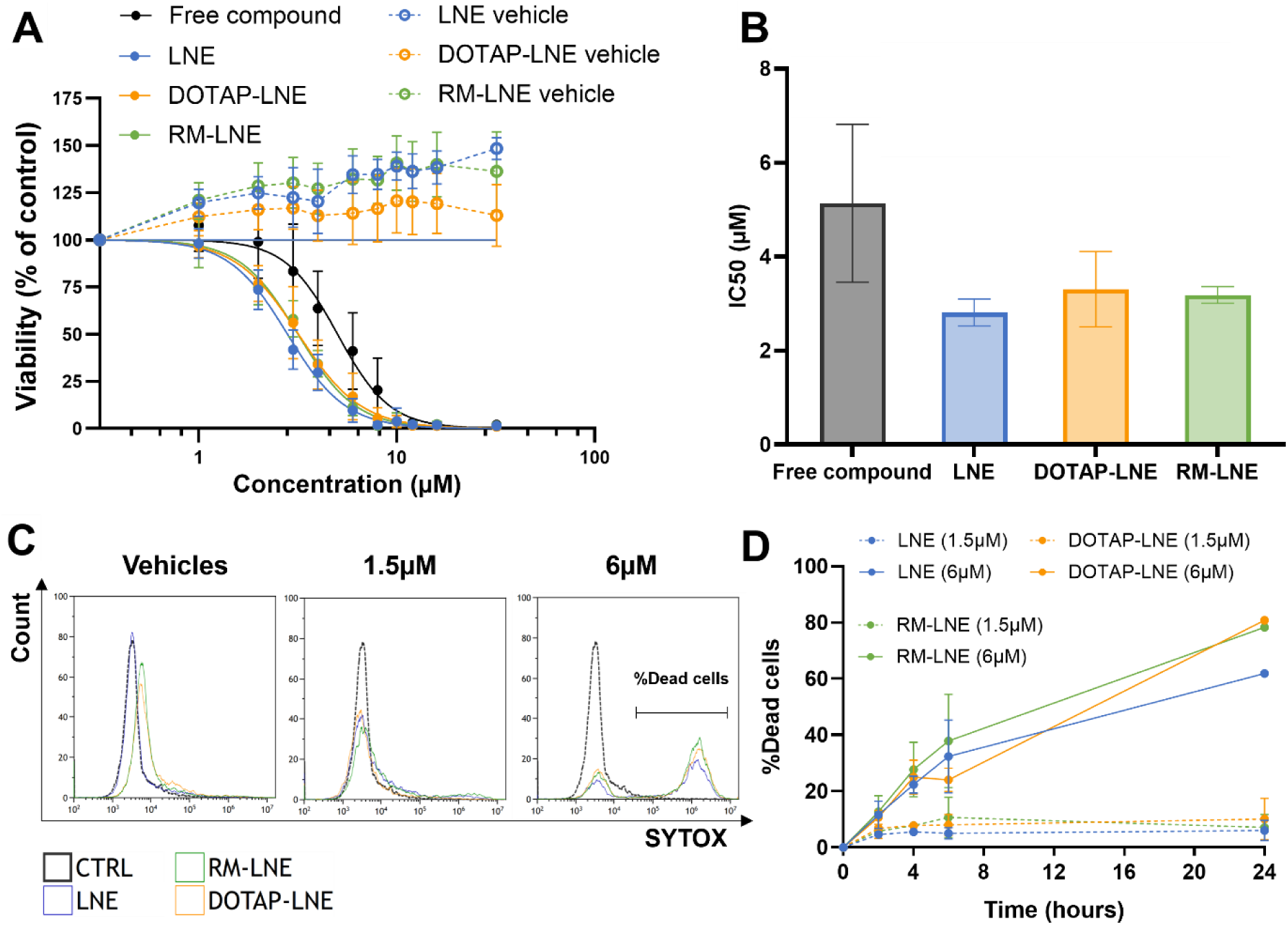
In vitro evaluation of cytotoxicity against H358 cells for ABD0171 encapsulated in LNE, DOTAP-LNE, and RM-LNE formulations. **(A)** Inhibition curves for formulations with encapsulated ABD0171, tested at concentrations ranging from 1 to 32 µM and compared to their respective vehicles (without ABD0171). Data are presented as mean ± SD of 15 replicates (3 independent experiments, 5 biological replicates each). **(B)** IC50 values for ABD0171 encapsulated formulations, based on the mean of three independent experiments (mean ± SD). **(C)** SYTOX low cytometry histograms comparing the untreated control to samples treated for 4 h with formulation vehicles or ABD0171-loaded formulations at 1.5 µM and 6 µM. The bar indicates the dead cell population. **(D)** Time-dependent evolution of dead cell populations following treatment with ABD0171-loaded formulations, based on data from (C).

To assess the safety profiles of the formulations, IC50 tests were also conducted using empty vehicles (nanoemulsions without the active compound). Interestingly, the viability curves (**Figure 7A**) for the empty formulations revealed cell viability exceeding 100%, suggesting that lipid-based nanoemulsions could enhance cell metabolism. This phenomenon can be attributed to the unique properties of lipid nanoparticles, which are known to interact with cellular membranes and enrich them with lipid substrates that can be utilized for energy production and biosynthetic pathways, as previously reported (42,43). Additionally, lipid nanoparticles have been shown to modulate metabolic activity by providing essential components for membrane synthesis and energy production, particularly in metabolically active cells such as cancer cells (44). These findings align with our observations, emphasizing that lipid-based nanoemulsions, in the absence of an encapsulated active pharmaceutical ingredient, can influence cellular behavior through metabolic support mechanisms. Interestingly, this effect may enhance the therapeutic efficacy of ALDH-targeted treatments by increasing cellular reliance on detoxification pathways such as the ALDH system. Consequently, cells with elevated metabolic activity due to lipid feed may be more susceptible to ALDH inhibition, thereby amplifying the cytotoxic action of ABD0171. However, this metabolic stimulation also warrants caution, as chronic exposure to LNCs has been associated with cellular stress and toxicity in other models (45,46), underscoring the need to carefully evaluate carrier-induced metabolic alterations.

Despite the presence of a positive charge on the surface of the DOTAP-containing formulations (DOTAP-LNE and RM-LNE), the observed lack of significant toxicity *in vitro* for the formulations without the drug (**Figure 7A**) suggested that a nanoparticle positive charge does not necessarily result in cellular toxicity. This finding aligned with the existing literature, which highlights that while positive charges on nanoparticle surfaces can enhance cellular uptake, they do not automatically lead to toxic effects. For example, Sharma et al. (41) indicated that while cationic nanoparticles may activate the complement system and induce toxicity, these effects were influenced by factors such as particle composition, biodegradability, and dose rather than being driven only by surface charge. Similarly, Goldstein et al. (40) emphasized that nanoparticle charge alone was not always predictive of drug efficacy or toxicity; instead, the interactions between the drug, delivery system, and cellular environment were pivotal in determining overall outcomes. Furthermore, studies on biodegradable nanoparticles have shown that adjusting surface properties, including charge, could reduce systemic toxicity by improving nanoparticle clearance, enhancing uptake by targeted cells, and minimizing immune system activation. In this regard, Salatin et al. (47) reported that surface charge, size, and shape are critical factors affecting both cellular uptake and the reduction of toxic effects. Despite their cationic nature, our lipid-based formulations benefited from biocompatible lipid components and optimized design, which included biodegradable properties, thus minimizing toxicity. As a result, these formulations did not induce significant adverse cytotoxicity *in vitro*, supporting their potential as safe and effective drug delivery systems.

### *In vivo* assessment of toxicity and efficacy using a chicken ChorioAllantoic Membrane (CAM) model

An *in vivo* toxicity and efficacy study of LNE formulations was conducted using a chicken CAM model grafted with tumors derived from the H358 cell line. The chicken CAM assay has emerged as a valuable intermediate model in cancer nanomedicine, bridging the gap between *in vitro* screening and *in vivo* rodent studies. Its rich vascularization and ability to support tumor grafts makes it particularly valuable for assessing drug distribution, tumor penetration, and early therapeutic response in a physiologically environment (48). Widely used to evaluate various drug delivery systems, including lipid nanoparticles, liposomes, and other formulations, the CAM model offers an ethical and cost-effective platform for preclinical screening, helping to support the rational selection of promising formulations prior to further validation in animal models (48,49).

In this study, this model was used for the simultaneous evaluation of compound toxicity and therapeutic potential, leveraging its ability to provide insights into tumor weight reduction and embryotoxicity. It was performed using four formulations of ABD0171 (ABD0171 free compound and ABD0171-loaded formulations LNE, DOTAP-LNE, and RM-LNE) at two drug dose levels (2.76 [1X] and 27.6 [10X] mg/kg). The high dose was administered once at Embryonic Development Day (EDD) 11, and the low dose was administered twice at EDD11 and EDD13. At a low dose (2.76 mg/kg [1X]), all formulations were well tolerated with embryotoxicity within the range of the NaCl 0.9% control group and no visible toxicity or CAM alterations (**Figure 8 second row** and **8B**). In contrast, at a high dose (27.6 mg/kg [10X]), high embryotoxicity was observed, especially for the control ABD0171 free compound, limiting sample availability (n = 6) for efficacy evaluation (**Figure 8B**). At high doses, formulations were better tolerated than the free compound, although they all caused severe CAM alterations, including hemorrhages, vessel coagulation, and blood clots **(Figure 8A first row),** necessitating the omission of a second administration. Interestingly, yellow oily droplets were observed on tumors treated with ABD0171-loaded DOTAP-LNE and RM-LNE at 27.6 mg/kg **(Supplementary Information Figure S7**), suggesting possible colloidal precipitation due to the CAM environment, which could induce localized release dynamics preventing the proper distribution of the ABD0171. Tumor weight analysis (**Figure 8C**) showed that at low dose, the ABD0171 free compound significantly reduced tumor weight by 25% compared to the untreated negative control. ABD0171 loaded DOTAP-LNE resulted in 28% tumor regression compared to the negative control, highlighting its therapeutic potential while maintaining a manageable toxicity profile. ABD0171 loaded RM-LNE also induced a decrease of tumor weight by 15%, although this effect was not significant compared with the untreated control.

**Figure 8:**
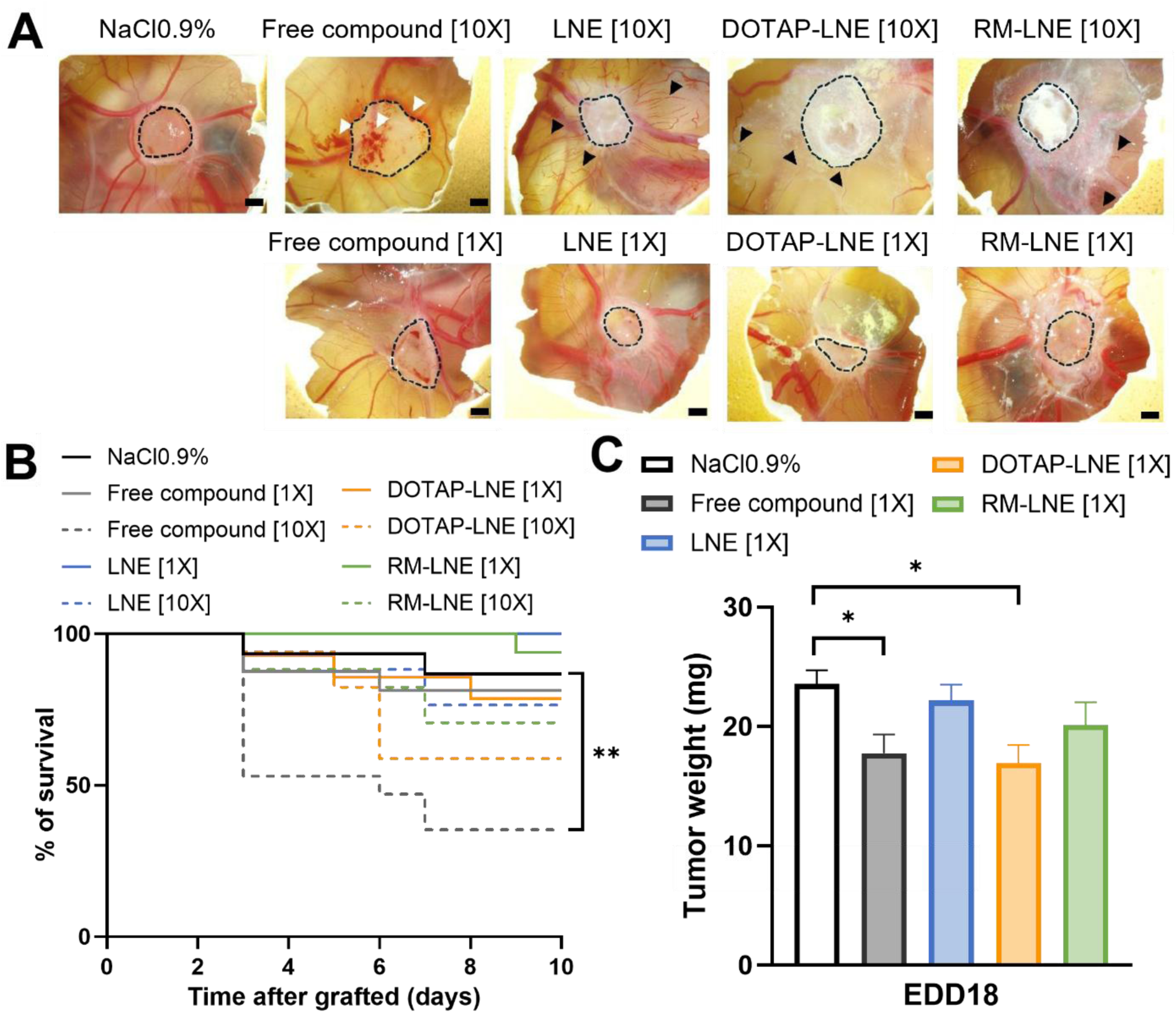
Evaluation of toxicity and therapeutic efficacy using the chicken CAM assay. **(A)** Images of the CAM 24 h after the first administration of each formulation for the two different doses ([1X] low dose of 2.76 mg/kg, [10X] high dose of 27.6 mg/kg). White arrows denoted hemorrhage, whereas black arrows indicated vessel coagulation and lysis. The tumor was outlined with a black dotted line; tumor boundaries were approximate owing to challenges in visualization when the CAM is altered. Scale bar = 2 mm. **(B)** Kaplan-Meier survival curves for groups treated with both doses. **(C)** Mean tumor weight (mg) measured across the experimental groups at the end of the study with low dose (2.76 mg/kg [1X]).

Thus, the formulation incorporating DOTAP as reverse micelles within the oily core led to a reduced tumor suppression efficacy compared to the one-pot formulation, where DOTAP is assumed to be predominantly localized at the surface. The physical state of the LNE oily core is known to be determined by the type of oil, its concentration, and the presence of stabilizing structures. Miglyol® 812N (caprylic/capric triglycerides) is a liquid at ambient temperature characterized by low viscosity and high fluidity, which facilitates the rapid release of encapsulated molecules (18). However, the incorporation of structures, such as DOTAP-based RM, can alter this state. Interactions between DOTAP and triglycerides can increase the local viscosity and modify the crystallinity of the lipid matrix, thus affecting overall fluidity (50,51). For instance, AOT reverse micelles created semi-structured or semi-solid regions within a Labrafac WL 1349 oily core (52), which slowed the diffusion of active molecules, resulting in more controlled release (51,53). Furthermore, the addition of stabilizing structures can influence the migration of DOTAP from the oily core to the nanoemulsion surface. In simpler systems, such as the DOTAP-mediated lung-specific delivery LNPs described by Dilliard et al. (15), the low viscosity and fluidity of the lipids allowed for easy migration of DOTAP from the oily core to the nanoparticle surface. In contrast, in more complex and "rigid" systems, such as LNE, the increased viscosity and more ordered organization of the oily phase could hinder this migration, reducing the DOTAP concentration at the surface. In summary, the physical state of the oily core in our LNE formulation likely influenced DOTAP migration and drug release by slowing or modulating its diffusion, thus reducing the therapeutic performance of the resulting formulation in this model. However, this effect could be further modulated in more complex environments such as complete blood circulation.

### *In vivo* toxicity evaluation in mice

To support the toxicity results obtained in the CAM model and prepare in depth efficacy studies in murine models, the maximum tolerated dose of the DOTAP-LNE formulation was evaluated in mice. In the CAM assay, delivering a therapeutically relevant dose of ABD0171 (27.6 mg/kg) required a high lipid concentration (∼210 mg/mL), which led to colloidal instability and phase separation on the tumor surface. These effects were likely due to strong interactions between the lipid-rich, cationic formulation and the CAM membrane, limiting systemic circulation and efficacy. This limitation highlighted the need to explore higher dosing regimens *in vivo*, under conditions better suited to systemic administration. Thus, a starting dose of 40 mg/kg was selected for murine studies, representing a modest 1.4-fold increase over the CAM dose. Such an adjustment is well within the range typically observed when translating CAM data to rodent models, where dose increases of 2–10× are often required due to interspecies differences in drug distribution and metabolism (54). The *in vivo* toxicity profiles of the DOTAP-LNE formulation showed a dose-dependent toxicity. At doses ≥ 80 mg/kg, acute respiratory distress and mortality occurred within 24 h post-administration. Necropsy revealed severe pulmonary edema, characterized by uniform red/purple discoloration and fluid accumulation in the lungs (**Supplementary information Figure S8)**, indicating direct pulmonary toxicity. This effect may be linked to the presence of the cationic lipid DOTAP, which has been shown to disrupt pulmonary surfactant and induce dose-dependent inflammation and tissue injury in murine lungs (55,56). Its strong pulmonary tropism (57,58) may further contribute to localized toxicity. However, a contribution from the active compound itself cannot be excluded, as high systemic doses of ABD0171 may also trigger off-target effects. Regulatory toxicology data obtained with a compound structurally analogous to ABD0171 (data not shown), demonstrated that the ALDHin are metabolized through hydrolysis of the thioester group of the molecule by esterases (**Figure 5A**), leading to the release of the methyl mercaptan gas. As a result, rodents, known for their high esterase activity and low pulmonary volume, are particularly sensitive to pulmonary distress following the administration of ALDHin. In particular, such pulmonary effects were not observed in dogs, demonstrating species-dependent toxicity profiles. The observed toxicity is therefore likely the result of a combined effect of the cationic carrier and the pharmacological agent, underscoring the importance of optimizing both the formulation composition and the dosing regimen. When the dose and the frequency of DOTAP-LNE administration was reduced (40 mg/kg, twice weekly over four weeks), tolerability significantly improved, with no observed respiratory distress or mortality (**Figure S9 in Supplementary Information**). This optimized protocol balanced safety with therapeutic dosing, highlighting the importance of adjusting treatment regimens for cationic lipid-based nanosystems for the delivery of cytotoxic compounds.

Taken together, our results demonstrate that the therapeutic benefits of ALDHin formulated as cationic lipid nanoemulsions necessitate careful optimization of dosing regimens and administration frequencies to mitigate off-target toxicities while maintaining therapeutic efficacy.

## CONCLUSION

In this study, we successfully developed, optimized, and characterized lipid nanoemulsions (LNEs) incorporating DOTAP, a cationic lipid known for its potential for targeted pulmonary drug delivery. DOTAP incorporating LNE were designed with and without reverse micelle structures, for the delivery of a second-generation ALDH inhibitor, ABD0171. Using fluorescence spectroscopy and dynamic light scattering (DLS), we demonstrated the feasibility of forming reverse micelles (RMs) within the oily core of LNEs and validated their ability to encapsulate DOTAP. We successfully encapsulated the active molecule ABD0171, with a high encapsulation efficiency (100%) and stability, consistent with its elevated lipophilicity and compatibility with the lipid core. Structural analyses revealed that incorporating DOTAP into the LNE systems increased the particle size and influenced the curvature parameters without compromising the stability. Positive zeta potential values were consistently observed across formulations, reflecting high positive surface charge and allowing colloidal stability of these nanoemulsions during long-term storage.

*In vitro*, the IC50 values of all ABD0171 loaded formulations were consistently below 5 µM, indicating strong therapeutic potential against H358 NSCLC cells. Moreover, despite the presence of cationic charges in DOTAP-containing formulations, no significant toxicity was observed in the absence of ABD0171, highlighting the biocompatibility of the formulations. However, in the CAM model, signs of local toxicity at high doses were observed for all formulations, regardless of composition. *In vivo* evaluation in mice further revealed dose-dependent pulmonary toxicity for the DOTAP-LNE formulations, highlighting the need for careful dose optimization to mitigate the adverse effects associated with cationic lipids for the delivery of ABD0171, especially in sensitive species. At lower doses, the DOTAP-containing LNE formulations showed therapeutic efficacy with manageable toxicity in the chicken CAM assay, demonstrating the possibility of fine-tuning the dosage and administration protocols for maximizing the therapeutic benefit of these formulations.

This work highlights the potential of reverse micelle-based LNEs for encapsulating lipophilic drugs along with DOTAP, known to improve lung tropism of lipid formulations (15). Future studies will explore the use of reverse micelles to encapsulate as well hydrophilic molecules for combination therapy and further optimize LNE formulations to improve lung cancer therapy, by promoting drug delivery efficiency along with reduced toxicity. Overall, this study positioned DOTAP-containing LNEs for ALDHin delivery as promising platforms in lung cancer therapy. These nanoemulsions have the potential to enter clinical trials and offer significant benefits to patients with NSCLC.

## ASSOCIATED CONTENT

Supporting information

### Author contributions

Maria Irujo: Conceptualization, methodology, investigation, formal analysis, data curation, visualization, writing, review and editing.

Laura Poussereau: Investigation (formulations and *in vitro*), data curation, writing. Maïssam Ezziani: Investigation (formulation and *in vitro*), data curation.

Clotilde Joubert, Chloé Prunier: Investigation (*in ovo* CAM model experiments), analysis.

Julien Vollaire, Véronique Josserand: Investigation (*in vivo* mice experiments), analysis.

Cédric Sarazin, Axel Kattar: Investigation (SD-TDA measurements), validation, resources.

Daphna Fenel, Eleftherios Zarkadas: Investigation (cryo-TEM imaging), resources.

Alice Gaudin: Supervision, writing, review and editing.

Mileidys Perez: Supervision, writing, review and editing.

Isabelle Texier: Supervision, project administration, funding acquisition, writing, review and editing.

## ACKNOWLEDGEMENTS

This work used the platforms of the Grenoble Instruct-ERIC centre (ISBG; UAR 3518 CNRS-CEA-UGA-EMBL) within the Grenoble Partnership for Structural Biology (PSB), supported by FRISBI (ANR-10-INBS-0005-02) and GRAL, financed within the University Grenoble Alpes graduate school (Ecoles Universitaires de Recherche) CBH-EUR-GS (ANR-17-EURE-0003). The IBS-ISBG EM facility is supported by the Auvergne-Rhône-Alpes Region, the Fondation Recherche Medicale (FRM), the fonds FEDER and the GIS-Infrastructures en Biologie Sante et Agronomie (IBISA). We thank Guy Schoehn for establishing and managing the IBS-ISBG cryo-electron microscopy platform and for providing access, training and support.

This work was supported by the French National Research Agency (ANR) through Labex ARCANE/CBH-EUR-GS program (ANR-17-EURE-0003).

## DISCLOSURE STATEMENT

Mileidys Perez, Alice Gaudin and Maria Irujo are employees of Advanced BioDesign which owns and develops the ABD0171 inhibitor. Laura Poussereau and Maïssam Ezziani were interns at Advanced BioDesign at the time experiments were performed. Mileidy Perez is a shareholder and the Chief Scientific Officer of Advanced BioDesign. All the other authors declare no competing interest.

The authors used ChatGPT (OpenAI) to assist with language refinement. All scientific content and interpretation were generated and validated by the authors.

