## Supplementary information for "Lipid nanoemulsion incorporating DOTAP reverse micelles as clinically translatable carriers of ALDH inhibitors for lung cancer therapy"

### SUPPORTING INFORMATION

| Formulation | [Lipid] (mg/mL) | | Final lipid composition (mg) | | | | | $V_{RM}$ added (mg) | [ABD-0171] (mg/mL) |
|---|---|---|---|---|---|---|---|---|---|
| | Initial | Final | Mygliol | Kolliphor | Lipoïd | DMPE-PEG$_{2000}$ | DOTAP | | |
| LNE (vehicle) | 840 | 210 | 663 | 205 | 158 | 9 | / | / | / |
| DOTAP-LNE (vehicle) | 840 | 210 | 663 | 205 | 38 | 9 | 120 | / | / |
| RM-LNE (vehicle) | 252 | 210 | 663 | 205 | 38 | 9 | 120 | 735 | / |
| LNE | 840 | 210 | 663 | 205 | 158 | 9 | / | / | 16 |
| DOTAP-LNE | 840 | 210 | 663 | 205 | 38 | 9 | 120 | / | 16 |
| RM-LNE | 252 | 210 | 663 | 205 | 38 | 9 | 120 | 735 | 16 |

**Table S1:** Mass composition and key parameters of formulations.

**Table S1** presents the mass composition of each formulation discussed in this study. The lipid concentration ([Lipid], mg/mL) corresponds to the mass of all excipients relative to the amount of water in the formulation. The amount of RM added in RM-LNE formulations is expressed as the mass proportion of RM relative to the initial LNE lipid mass. The mass composition reflects the amount of each component added to the formulation. The ABD0171 concentration corresponds to the concentration calculated using the amount of drug within the formulation over the total volume of formulation including excipients and water.

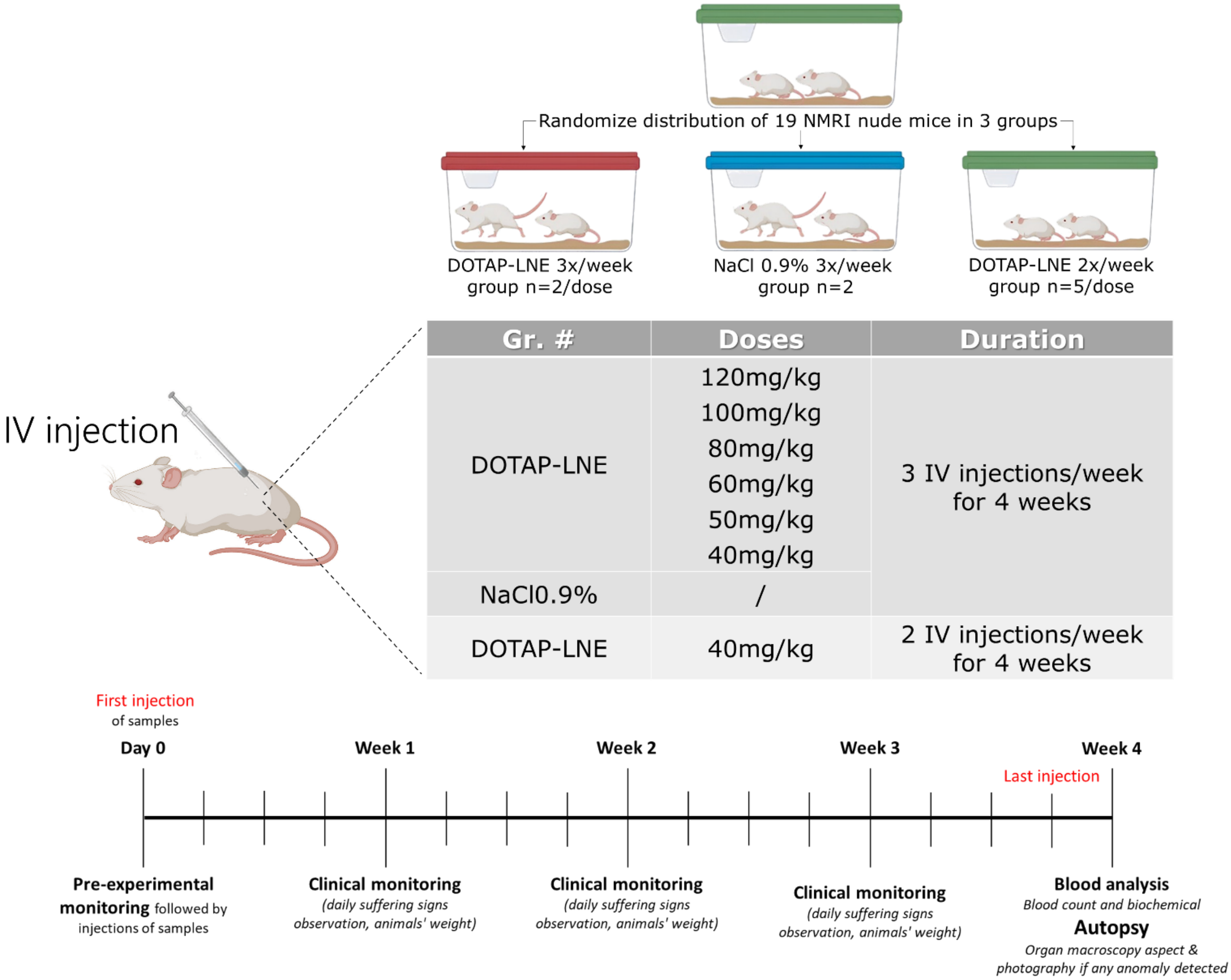


**Figure S1:** Experimental design for toxicity study in mice with different formulations to establish maximum tolerated dose and injection frequency for further efficacy studies.

Formulations were prepared by dilution in 0.9% NaCl and administered via tail vein intravenous bolus injections (8 µL/g). To determine the maximum tolerated dose, body weight, behavior, and distress were monitored with euthanasia performed if ethical limits were exceeded. All the mice were euthanized for organ analysis at the end of the study.

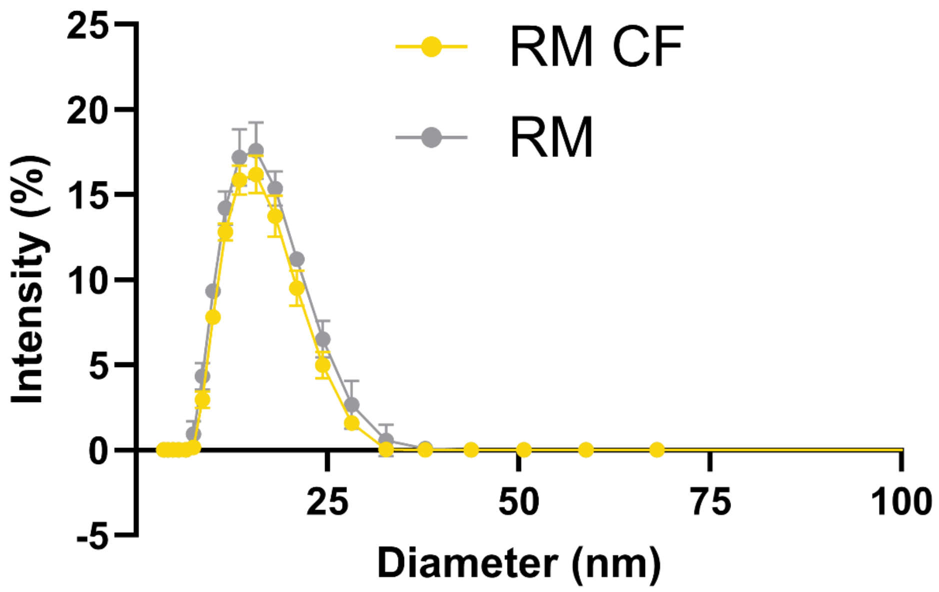


**Figure S2:** Size distribution and PDI of RM with and without CF.

**Figure S2** presents hydrodynamic diameter distribution of reverse micelles prepared with and without CF, measured using a Zetasizer Nano ZS (Malvern Instruments, UK). Dynamic light scattering (DLS) measurements for RM(CF) were performed under identical conditions as for RM without CF. As indicated by the homogeneous DLS distribution, all formulations exhibited a PDI $< 0.25$, reflecting high uniformity in micelle size.

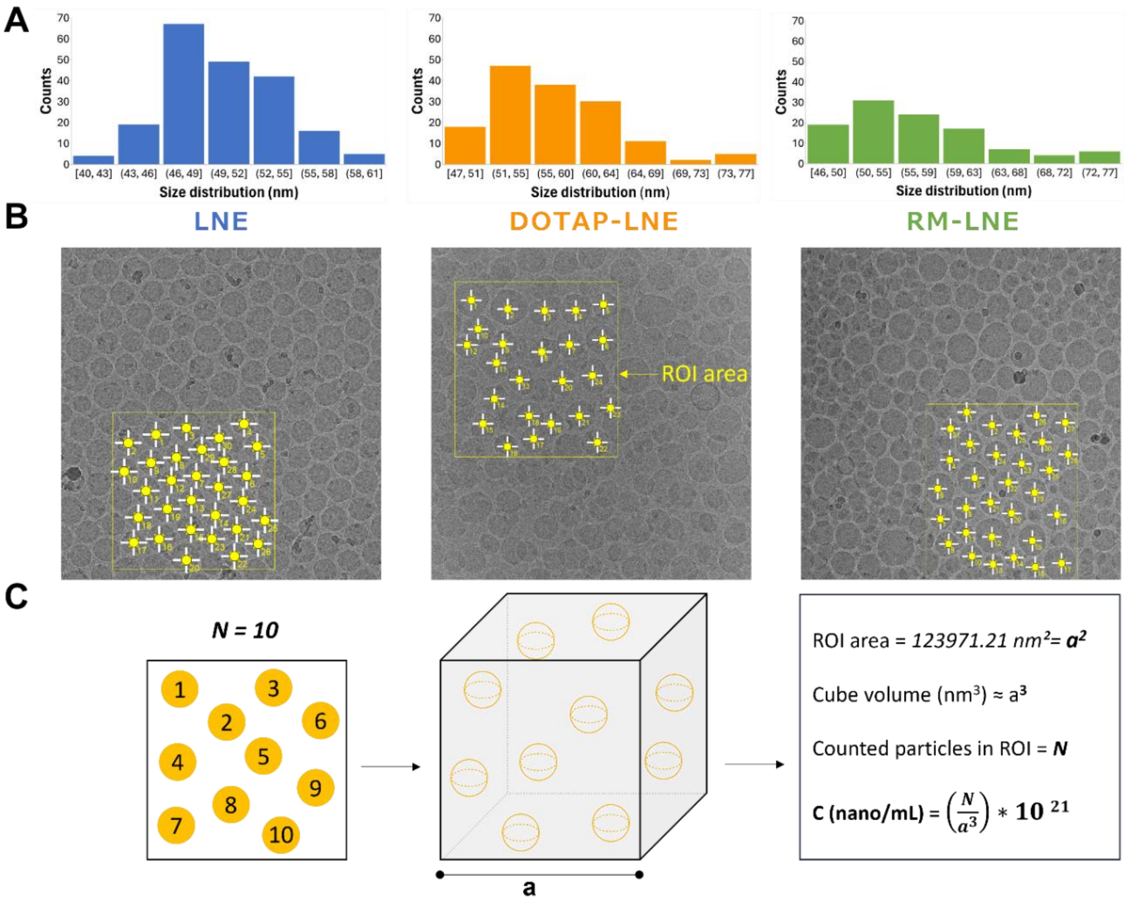


**Figure S3:** (A) Size distribution histograms obtained from cryo-TEM images using ImageJ for the different formulations. (B) Representative cryo-TEM images showing the regions of interest (ROI) used for nanoparticle counting. (C) Schematic illustrating the method used to estimate nanoemulsion concentration (nano/mL). Generated data are the mean of 3 images with 73kX resolution for each formulation.

To estimate the concentration of nanoparticles (nano/mL) in each formulation, cryo-TEM images were analyzed using the ImageJ software to count individual nanoparticles within a defined square region of interest (ROI). The ROI area was extracted from the software and used to calculate the square dimension $\boldsymbol{a}$. Reconstructing a theoretical cubic volume and assuming a homogeneous distribution of particles in the sample, the number of particles counted in 2D (N) was extrapolated to a 3D cube of volume $\boldsymbol{a^3}$, where $\boldsymbol{a}\,(\boldsymbol{nm}) = \sqrt[2]{ROI\ area}$. The resulting nanoemulsion concentration was then calculated using the formula ($10^{21}$ being the converting factor between $nm^3$ and mL):

$$C\ (nano\ /\ mL) = \frac{N}{a^3} * 10^{21}$$

This approach provides an order-of-magnitude estimate of nanoparticle concentration based exclusively on cryo-TEM data, without requiring external calibration methods.

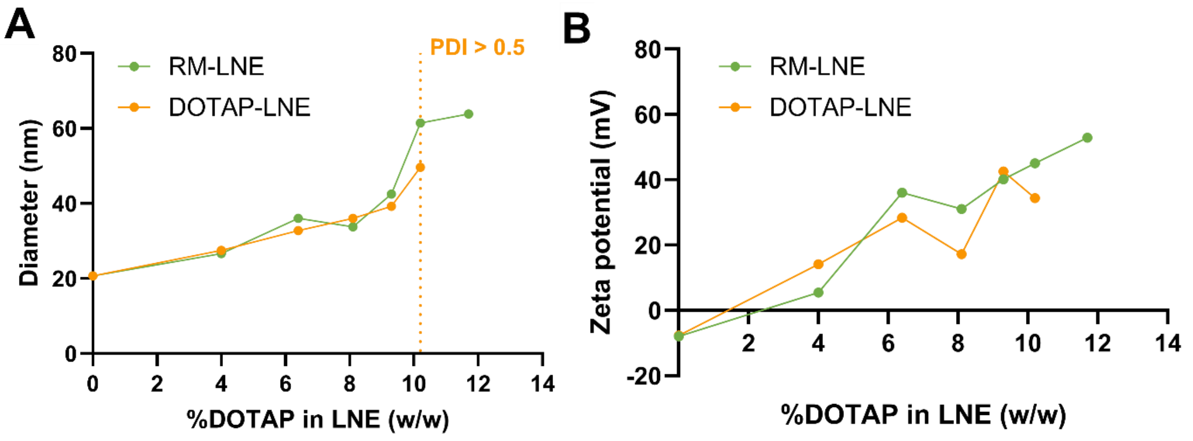


**Figure S4:** Optimization of DOTAP incorporation in RM-LNE and DOTAP-LNE formulations: Size and zeta potential analysis.

After fixing the DOTAP and water quantities in the RM preparation process, the study aimed to identify the optimal RM amount required to maximize encapsulation while ensuring a homogeneous, stable, and easily realizable formulation. Concurrently, the optimal DOTAP quantity was also evaluated using the one-pot formulation process, in which DOTAP was mixed with other components prior to thermocycling to yield DOTAP-LNE. Nanoparticle diameter and zeta potential were measured when varying the DOTAP quantity in both RM-LNE and DOTAP-LNE nanosystems. The diameter (Figure S4A) and zeta potential (Figure S4B) were plotted against the percentage of DOTAP relative to the total lipid mass in the formulations.

Figure **S4A** illustrates a steady increase in particle size for both RM-LNE and DOTAP-LNE formulations with increasing DOTAP concentration. However, beyond 10.2% DOTAP, the DOTAP-LNE formulations became heterogeneous (PDI > 0.5), indicating a limit of structural compatibility. In contrast, RM-LNE remained homogeneous and stable (PDI < 0.2) up to 11.7% DOTAP, suggesting that embedding DOTAP within reverse micelles allowed for higher incorporation without disrupting nanoemulsion stability. This enhanced capacity was likely due to a larger volume that can be occupied by DOTAP within the oily core before saturating the nanoemulsion surface.

In **Figure S4B**, zeta potential also increased with DOTAP concentration for both formulations, indicating progressive enrichment of the nanoemulsion surface with cationic lipid. For DOTAP-LNE, the zeta potential increased sharply at low DOTAP concentrations, then showed a slower rise beyond ~6%, suggesting a progressive saturation of the particle surface as the formulation approached its structural limit. At higher DOTAP content (≥ 10.2%), the increase became less consistent, reflecting the onset of formulation instability. A similar trend was observed for RM-LNE, although the zeta potential increased more gradually over the entire range, which may reflect a slower redistribution of DOTAP from the core to the interface.

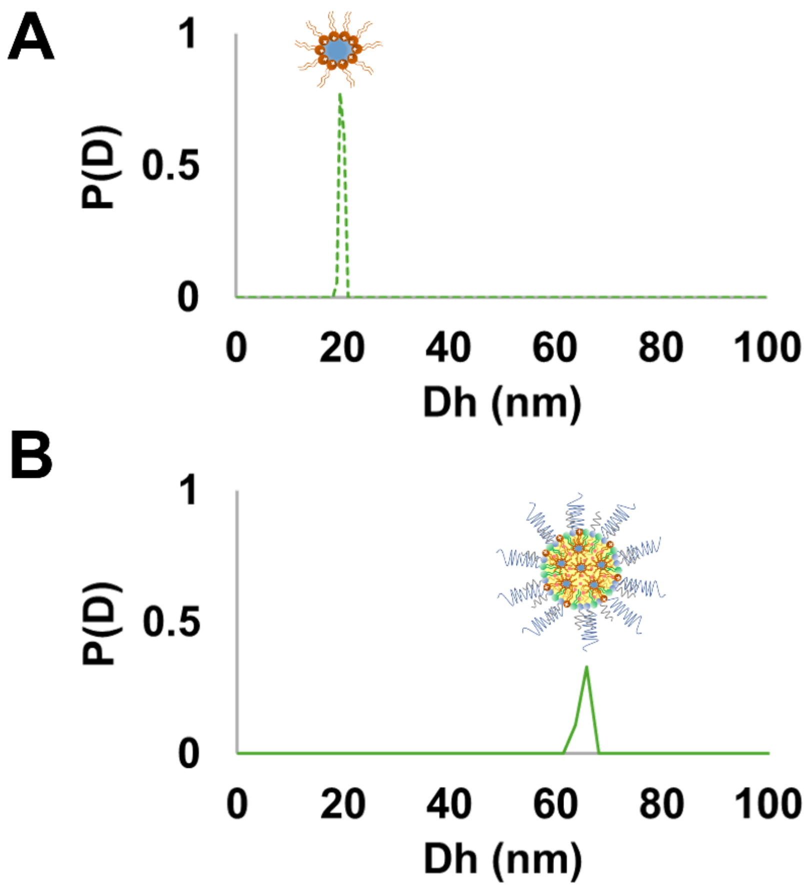


**Figure S5:** SD-TDA analysis of free DOTAP RM (A) and RM-LNE formulation (B) measured at 200 nm.

To assess the structural incorporation of reverse micelles (RM) into the lipid core, SD-TDA measurements were performed at 200 nm, corresponding to the specific absorbance of DOTAP. Free DOTAP RM exhibited a narrow distribution centered at ~14 nm (A), while the RM-LNE formulation showed a monodisperse peak at ~55 nm (B). The absence of any second population at ~14 nm confirms the successful encapsulation of RM within the lipid nanoemulsion droplets.

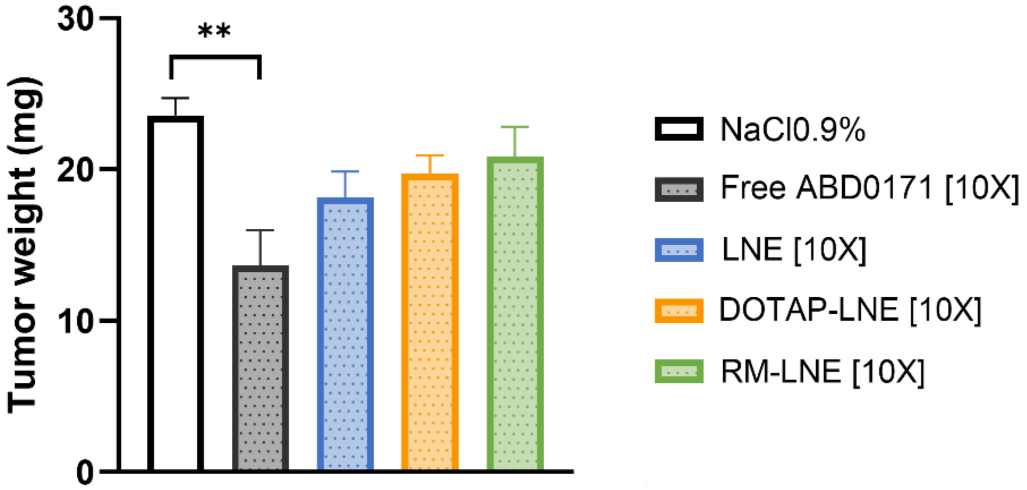


**Figure S6:** Mean tumor weight (mg) measured across experimental groups at the end of the study with [10X] dose.

The free ABD0171 group exhibited a significant antitumor effect, with a 42% reduction in mean tumor weight compared to control. However, this analysis was performed on a reduced number of samples (n = 6) due to the toxicity observed in this group. The formulations LNE, DOTAP-LNE, and RM-LNE, while better tolerated, did not induce statistically significant tumor weight reductions at this dose. This discrepancy may result from the lower effective dose delivered due to formulation-related factors such as lipid overload, altered biodistribution or precipitation at the CAM surface, as shown in Figure S7. It is important to note that no dedicated vehicle only control group (formulation without active compound) was included at this dose, which limits the interpretation of lipid-related effects independently of the drug.

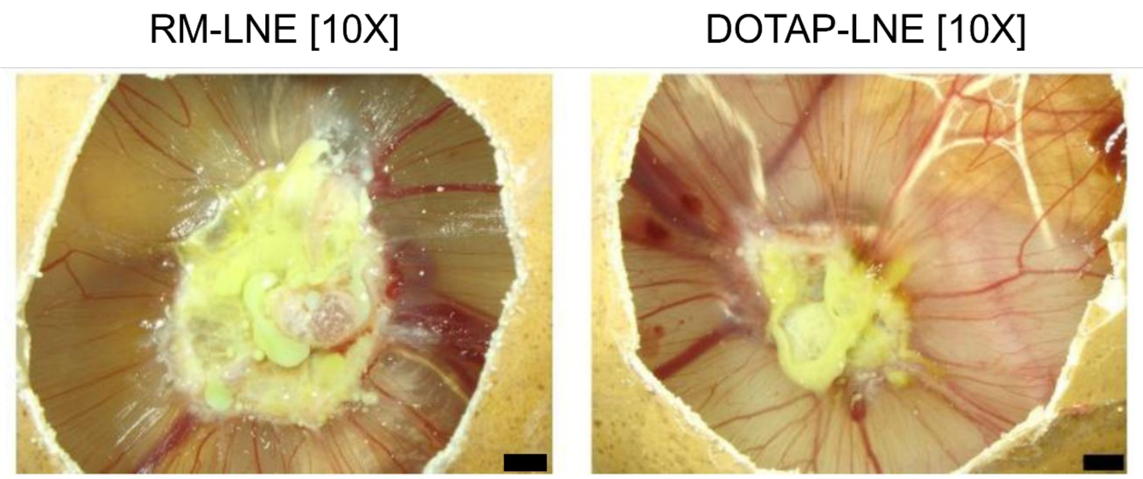


**Figure S7:** Yellow deposition on the chorioallantoic membrane (CAM) tumors at 27.6 [10X] mg/kg of ABD0171 formulation

Figure S7 illustrates the yellow deposition observed on tumors over the CAM at the end of the study (EDD18) in embryos treated with high doses (27.6 mg/kg) of RM-LNE and DOTAP-LNE. This deposition was not observed in the low dose treatment groups. The figure includes representative images of the CAM at EDD18, with a scale bar of 2 mm.

The administered dose of 27.6 mg/kg was achieved using formulations with an ABD0171 concentration of approximately 18 mg/mL. At this concentration, the lipid content reached 210 mg/mL (see Table S1). The high lipid content interacted directly with the biological components of the CAM, adhering to its surface. This interaction, combined with the positive charge of the formulations, caused colloidal instability, leading to phase separation and precipitation of the nanosystem on the tumor surface.

These results suggest that achieving the delivery of high ABD0171 doses requires increasing the drug loading within the oil core of the formulations. This approach would allow reducing the lipid excipient dose, improving colloidal formulation stability and facilitating systemic circulation within the embryo's bloodstream, thereby enhancing the therapeutic efficacy *in ovo* at elevated ALDHin doses.

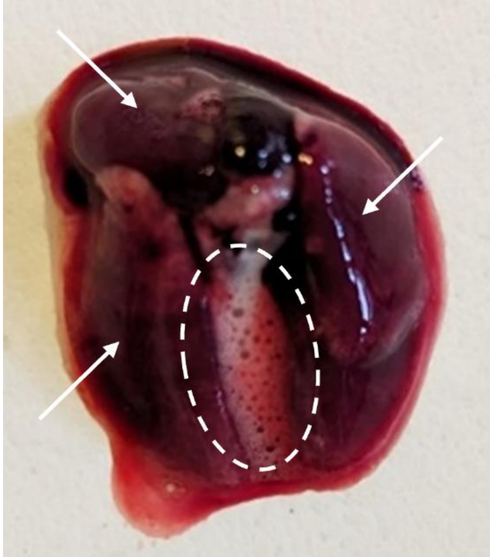

**Figure S8:** *Ex vivo* lung image obtained following injections at doses ≥ 80 mg/kg of DOTAP-LNE formulation.

Figure S8 illustrates necropsy findings from animals that died overnight following injections of ≥ 80 mg/kg of DOTAP-LNE formulation, highlighting severe pulmonary damage characterized by an edematous phenotype. The *ex vivo* image reveals a uniform red/purple discoloration of the lungs, marked by white arrows, and the presence of mossy fluid, delineated with a dashed white circle.

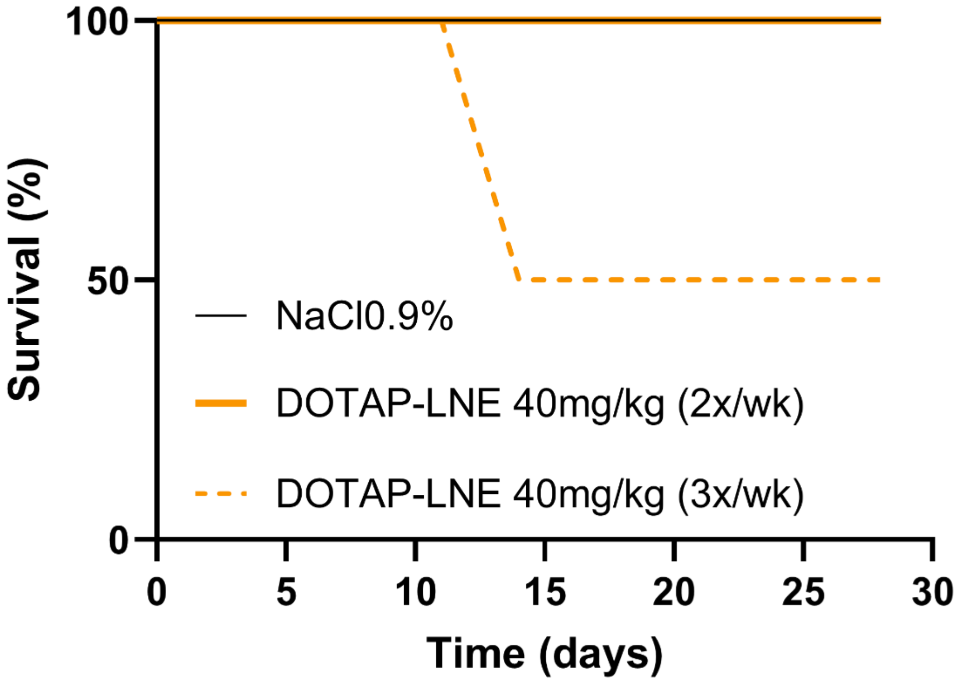


**Figure S9:** Kaplan-Meyer survival analysis for toxicity assessment of DOTAP-LNE in NMRI nude healthy mice.

Figure S9 presents the Kaplan-Meier survival curves of a toxicity study conducted in NMRI nude mice (n = 2-5 animals per group). The study compared the survival of mice treated with ABD0171 loaded DOTAP-LNE administered intravenously (IV) at a dose of 40 mg/kg, three times (dotted orange line) or twice (solid orange line) per week for four weeks.

Administration three times per week led to a 50% survival rate by the midpoint of the treatment period, indicating lower tolerability of this formulation under these dosing conditions in healthy mice. However, when the frequency of DOTAP-LNE treatment was reduced to twice per week (dotted orange line), survival improved to 100% by the end of the four-week treatment period.

These results highlight the importance of optimizing treatment frequency to mitigate toxicity in healthy subjects when using the DOTAP-LNE formulation.
